# A single strand-based library preparation method for unbiased virome characterization

**DOI:** 10.1101/2024.03.31.587488

**Authors:** Xichuan Zhai, Alex Gobbi, Witold Kot, Lukasz Krych, Dennis Sandris Nielsen, Ling Deng

**Affiliations:** Section for Food Microbiology, Gut Health and Fermentation, Department of Food Science, University of Copenhagen, Rolighedsvej 26, 1958 Frederiksberg C, Denmark; Section of Microbial Ecology and Biotechnology, Department of Plant and Environmental Sciences, University of Copenhagen, Thorvaldsensvej 40, 1871 Frederiksberg C, Denmark

**Author notes:** Current address: European Food Safety Agency (EFSA), Via Carlo Magno 1A, 43126 Parma, Italy.

**Keywords:** single-stranded library, phage mock community, ssDNA virome, dsDNA virome, RNA virome, gut virome, modification

## Abstract

The gut virome is an integral component of the gut microbiome, playing a crucial role in maintaining gut health. However, accurately depicting the entire gut virome is challenging due to the inherent limitations and biases associated with current sequencing library preparation methods. To overcome these problems, we repurposed the ligation-based single-stranded library (SSLR) preparation method for virome studies. We demonstrate that the SSLR method exhibits exceptional efficiency in quantifying viral DNA genomes (both dsDNA and ssDNA) and outperforms existing double-stranded (Nextera) and single-stranded (xGen, MDA+Nextera) library preparation approaches in terms of minimal amplification bias, evenness of coverage, and integrity of assembling viral genomes. The SSLR method can be utilized for the simultaneous library preparation of both DNA and RNA viral genomes. Furthermore, the SSLR method showed its ability to capture highly modified phage genomes, which were often lost using other library preparation approaches.

## Main

The gut microbiome constitutes a diverse array of microbes, comprising bacteria, archaea, viruses/bacteriophages, and fungi, residing in the human intestine ^1^. Bacteriophages (or phages for short), among them, are believed to be highly abundant constituents of the human microbiome possibly equal in numbers to bacteria. They play important roles in modulating the diversity and abundance of gut bacteria, maintaining a dynamic equilibrium ^1, 2^. There is growing evidence linking gut virome dysbiosis to human diseases such as inflammatory bowel disease ^3–5^, severe acute malnutrition ^6^, alcoholic liver disease ^7^, type 2 diabetes ^8^ stunting ^9^ and recently also asthma ^10^. Studies have also demonstrated that in fecal microbiota transplantation to treat e.g. recurrent *Clostridioides difficile* (rCdiff) the virome component is important for treatment efficacy and even that sterile filtered feces used for so-called fecal filtrate transplantation is able to cure rCdiff ^11–13^. Consequently, there is growing interest in studying the gut virome to understand its role in various diseases and its potential clinical applications.

However, this effort is hampered by persistent biases and inconsistencies across studies due to non-standardized and non-optimized pipelines. Every stage of the virome exploration process, from sample collection to bioinformatic analysis, is crucial, with virome characterization of complex microbial communities relying on essential steps such as virus-like particle isolation/purification, viral DNA/RNA purification, and sequencing library preparation and shotgun high-throughput sequencing ^14–17^. Furthermore, despite advances during recent years, virus identification is still challenging due to their variability, small genome sizes, and genetic mosaicism. Further, virome studies (distinct from bacteriome studies) present a unique challenge with diverse genome types (dsDNA, ssDNA, dsRNA, and ssRNA) and different topologies (linear, circular, or fragments) ^18, 19^, complicating library preparation and down-stream analysis ^1, 3, 20^, emphasizing the need for universal or standardized virome sequencing library construction methods for improved reproducibility and comparability across studies.

Commonly used library preparation methods, such as randomly amplified and linker-amplified shotgun libraries, have limitations, as these methods are restricted to target only dsDNA viruses and can result in uneven coverage of viral genomes, particularly when the viruses are very different in proportional abundances ^19, 21^. The transposon-based method, while fast and requiring low-input template, is still restricted to dsDNA ^22^. To overcome these limitations, multiple displacement amplification (MDA) has been utilized, increasing DNA amounts for low biomass virome samples and converting ssDNA into dsDNA for further library preparation. However, MDA distorts the ratios of different viral DNA forms by over-amplifying small circular ssDNA genomes and unevenly amplifying linear genomes ^23–25^. More recently, single-stranded based methods for library construction have been applied, such as the xGen ssDNA & Low-Input DNA library preparation kit from IDT (previous name Accel-NGS kit from Swift Biosciences). Though promising, the cost per sample is relatively high and it involves multiple purification steps lowering throughput.

Currently, RNA viruses, especially dsRNA viruses, are significantly undersampled due to the instability of RNA and its incompatibility with common DNA sequencing library preparations ^26, 27^. As a result, DNA and RNA viruses are analyzed separately ^28^. While recent studies have revealed a greater diversity of environmental RNA viruses than previously thought ^26, 29, 30^, little information is available on human gut RNA viral communities ^31^. The commonly used RNA-Seq technology for investigating RNA viruses requires large amounts of sample material, the preparation process is expensive and time-consuming, and can be further complicated by DNA contamination ^32–34^.

To address these challenges, we drew inspiration from the Single Reaction Single-stranded LibrarY (SRSLY) method, originally designed for the library preparation of cell-free DNA and oligo sequencing ^35^. We have extended and improved this method for gut virome sequencing (both DNA and RNA) with cost-effectiveness and timesaving in mind. We compared the capability and quantitative accuracy of SRSLY with three different virome library preparation methods (Nextera, MDA amplification and xGen) for Illumina short-read sequencing. We evaluated the types and levels of sequencing bias generated by these protocols using a wide range of DNA/RNA phage mock communities and performed qualitative and quantitative analyses using a diverse mock community with different ratios of 4 different genome types (dsDNA, ssDNA, dsRNA and ssRNA) to further verify the method’s applicability for virome studies. Additionally, we assessed reliability, including error rates, composition bias and assembling accuracy of these methods. Finally, we validated the utility and performance of the methods for human fecal virome analysis.

## Results

### SSLR is an effective method for quantifying DNA phage genomes

We conducted sequencing library preparation using three DNA mock communities (Mock A, B and C; **Fig. S1A** and **S1E**) containing different ratios of dsDNA to ssDNA phages (10%, 50%, and 90% of dsDNA, **Table S1**) with five different methods, namely, Nextera, MDA_0.5h, MDA_1.5h (0.5h and 1.5h referring to the amplification time), xGen and SSLR. Overall, all the phage genomes were detectable even when the total dsDNA inputs were as low as 0.20 ng in the high ssDNA genomes mock (Mock C, **Fig. 2A**). However, the efficiency of quantification varied between methods. Nextera library preparation significantly underestimated ssDNA phages (24 to 35 fold, **Fig. 2A** and **Table S1**), where only 7 of 9 genomes could be captured (**Table S2**), making it unsuitable for ssDNA genomes studies. Although MDA can be very helpful for ssDNA genome studies, we observed a selective amplification bias of ssDNAs (3-fold, **Table S1**) even with short-time amplification (MDA_0.5h). Notably, no significant amplification biases were observed in mocks with a high percentage of ssDNA genomes (≥ 50%, about 1-fold, **Table S1**). In contrast, both SSLR and xGen accurately recovered the percentage of ssDNA genomes when present in high ratios (≥ 50%, Mock B and C), but slightly underestimated ssDNA genomes when present in low ratio inputs (about 10%, Mock A, **Fig. S2A**).

**Fig. 1.**
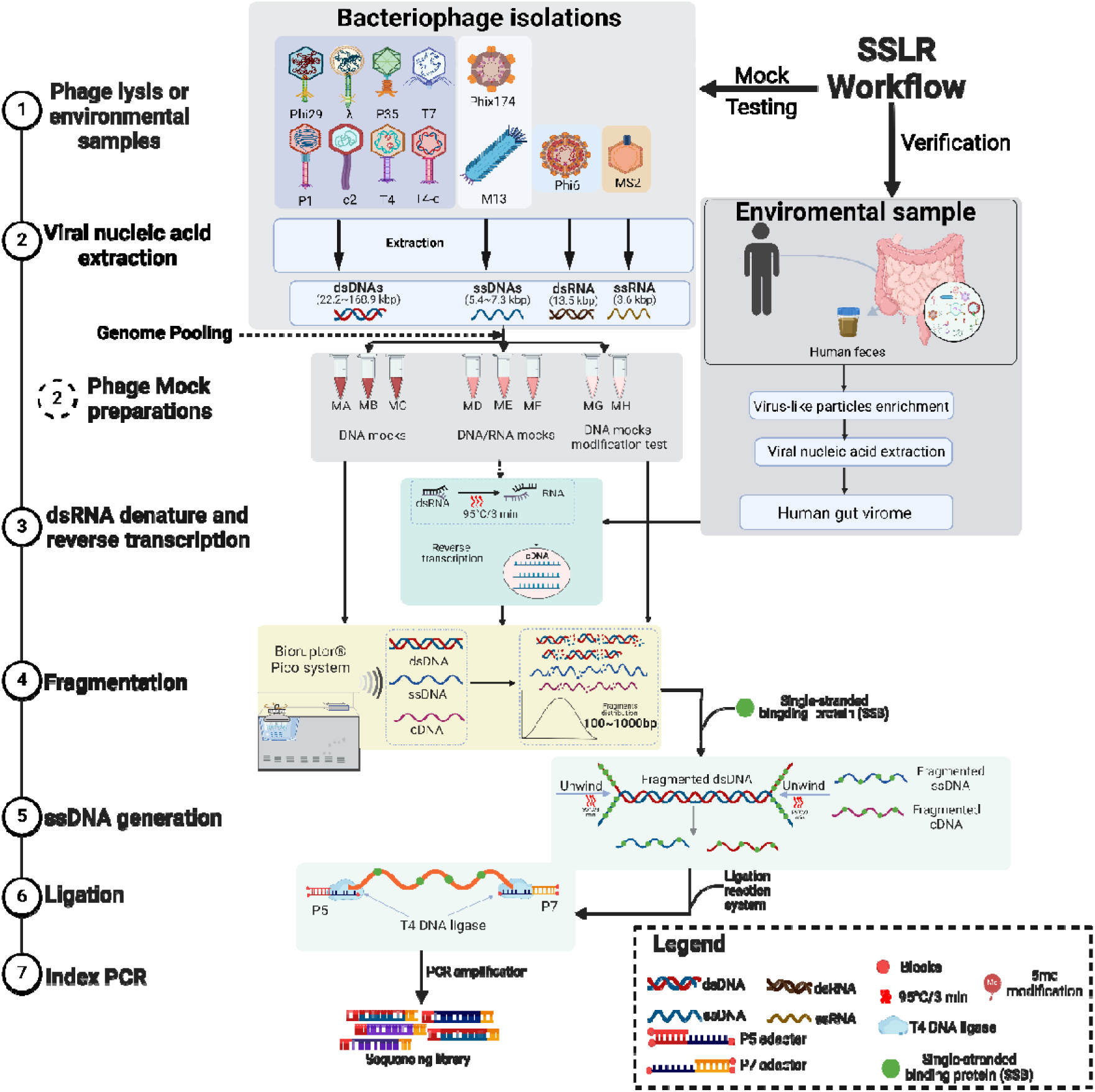
| Overview of study workflow. The reproposed library preparation method (SSLR) was indicated with 7 steps (detailed information can be found from **Fig. S1** and **Supplementary file1**). Five different libraries were used to prepare and sequence three artificial bacteriophage mocks containing different proportions of the ssDNA phages (phiX174 and M13mp18) mixed with the dsDNA phages. These phage genome abundance values were calculated based on the quantity of dsDNA and ssDNA phages measured Qubit dsDNA (or ssDNA) HS Assay kit. MA: Mock A with a ratio of ∼90:10 for dsDNA and ssDNA (**Fig. S1E**); MB: Mock B with a ratio of ∼50:50 for dsDNA and ssDNA; MC: Mock C with a ratio of ∼10:90 for dsDNA and ssDNA. MD: Mock D with a ratio of ∼90:10 for DNA and RNA; ME: Mock E with a ratio of ∼50:50 for DNA and RNA; MF: Mock F with a ratio of 10:90 for DNA and RNA (**Fig. S1F**). MG: Mock G contains high modification T4 genome (T4) with equal ratio of all genomes; MH: Mock H contains lower modification T4 genome (T4-c) with equal ratio of all genomes (**Fig. S1G**).

**Fig. 2.**
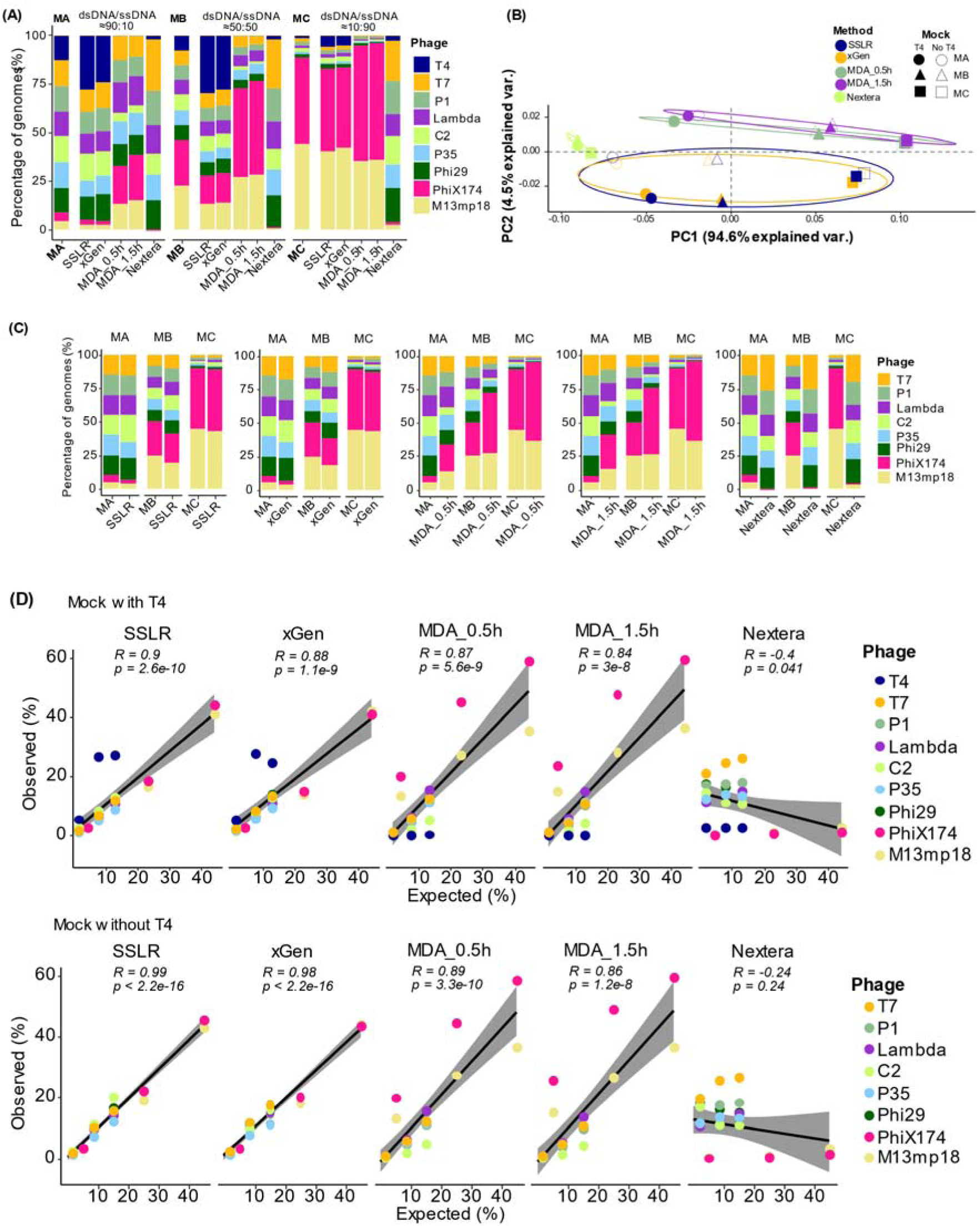
| Comparison of different library strategies for DNA mock communities. (A) Percentage of phage genomes with T4 generated by different library preparation methods. Different color indicate different phage genomes. The percentage of phage genomes was calculated by aligning the clean reads to customed database (**Supplementary file6**) with bowtie2 to assess the relative abundance of each DNA phage (**Table S1**). (B) Principal coordinates analysis (PCoA) plots of bray-Curtis distance matrices. PCoA was used to plot the beta diversity of mock-associated communities using the bray matrix. Different colors indicate different library preparation methods, different shapes indate different DNA mock communites. The dark-filled shapes display the mock with T4 genome and the non-filled without T4. For each axis, in square brackets, the percent of variation explained was reported. (C) Percentage of phage genomes without T4. (D) Pearson correlation coefficient (r) and two-tailed p-value between the expected and obtained read distributions (in percentage) in different DNAs phage mock communities with (top panel) or without (bottom panel) phage T4 genome. Five different libraries were used to prepare and sequence three artificial bacteriophage mocks containing different proportions of the ssDNA phage phiX174 and M13mp18 mixed with the dsDNA phage. These phage genome abundance values were calculated based on the quantity of dsDNA and ssDNA phages measured Qubit dsDNA (or ssDNA) HS Assay kit. MA: Mock A with a ratio of ∼90:10 for dsDNA and ssDNA; MB: Mock B with a ratio of ∼50:50 for dsDNA and ssDNA; MC: Mock C with a ratio of ∼10:90 for dsDNA and ssDNA.

Interestingly, we observed quite different quantifications of the T4 genome prepared with different methods (**Fig. 2A, 2B, 2D** and **Table S1**). Less than 2% of T4 genomes were sequenced in all the MDA-related libraries while 23% to 34% were reached in the Nextera libraries compared to their corresponding inputs. In contrast, both SSLR and xGen overestimated T4 genomes (1.8 to 3.6 folds, we later found this is due to under-quantification of T4 DNA concentration using Qubit for mock construction, see below), especially in the mock with low T4 genome input (Mock C, **Fig. 2A**, **2D** and **Table S1**).

We further examined the quantification of viral genomes without considering T4 in the mocks (**Fig. 2C** and **2D**) and found that the percentage of each genome was consistently accurate across different mock communities when the libraries were constructed by SSLR or xGen. These methods showed strong linear correlations between expected and observed percentage distributions (*p* < 2.2e-16, **Fig. 2D**). Genome size and GC-content of each genome had minimal influence on these quantifications (**Fig. S2B** and **S2C**), and both methods provided a similar overview of DNA phage communities (**Fig. 2B**). While the variations could be minimized for the MDA-related methods without including the T4 genome, the quantifications of ssDNA (especially phiX174) were less consistent than the quantification of dsDNA genomes due to their much higher sequencing depths (**Fig. S3** and **Table S1**). The quantification of phages using the Nextera method appeared to be more consistent when the dsDNA phages were calculated exclusively (**Fig. 2B** and **2C**).

### SSLR has less bias and higher accuracy in sequencing DNA phages compared to existing methods

We then assessed the efficiency and accuracy of the library prepared using different methods with the presence of the T4 genome. The overall mapping rate to the mock genomes from SSLR method was comparable to xGen but higher than Nextera and MDA-related libraries (**Fig. S4A**). Furthermore, SSLR had a lower error rate for the entire mock community compared to Nextera and MDA methods, similar to the rate produced by xGen method (**Fig. S4B** and **S4C**).

When we analyzed individual genomes from SSLR and xGen, and observed mostly even coverage among the genomes, except for sharp spikes near the start and end of each genome (**Fig. S3**). However, the coverage of ssDNA genomes was significantly increased in MDA-related methods and largely neglected by the Nextera method (**Fig. S3** and **Fig. S4D**). Additionally, the effect of GC content on individual genome profiles (GC bias) varied among different library preparation methods. MDA-related and Nextera libraries showed a similar pattern with coverage increasing as GC content increased. However, coverage biases from SSLR and xGen were not apparently associated to their GC content (**Fig. S5**).

Finally, we evaluated the assembly accuracy and efficiency of the DNA phage genomes in the mock community prepared with different library methods. At a same sequencing depth, the assembly from the SSLR prepared library exhibited perfect quality, including the fully reconstructed phage T4 genome (**Table S3**). In contrast, the other library methods provided less integrity with large gaps in the reconstructed genomes. Moreover, the hits of assembled contigs to the reference genomes from the SSLR prepared library showed one contig per reference genome (except for phage P1, where SSLR led to 2 contigs for P1 genome, **Table S4**), where the other methods resulted wide range hits from 0 to 22.

### SSLR can quantify highly modified viral DNA

Notably, the T4 genome, in which all cytosines are modified to 5-hydroxymethylcytosine (5-HMC) and further glucosylated (glc-HMC) ^36^, showed a distinct coverage pattern when prepared with MDA-related methods. The sequencing depths at all genome positions were quite low regardless of the MDA time (**Fig. S3**). This finding prompted further investigation by introducing an unmodified C-sites of T4-c genome, which instead have amber mutations in dCTPase and dHMase genes^37^. Quantification of the T4 genome by Nanodrop showed significantly higher concentration (3.03 – 3.45 folds) than Qubit measurements of the same samples, while the less modified T4-c genome showed less differences (1.65-fold, **Fig. 3A**). MDA amplification of T4 genome resulted in relatively lower amount of product with shorter fragments compared to phage genomes P1, T7 and T4-c (**Fig. 3B** and **3C**). Equal amounts of phage genome (11.11% of each, quantified by Nanodrop) was used for preparation of Mock G (with T4) and Mock H (with T4-c), we observed that T4-c still yield lower percentage after sequencing (**Fig. S6**), but had higher ratios (about 2-fold) than T4 in Nextera, SSLR and xGen method (**Fig. S1C**, **Fig. S1G**, **Fig. 3D** and **Table S5**).

**Fig. 3.**
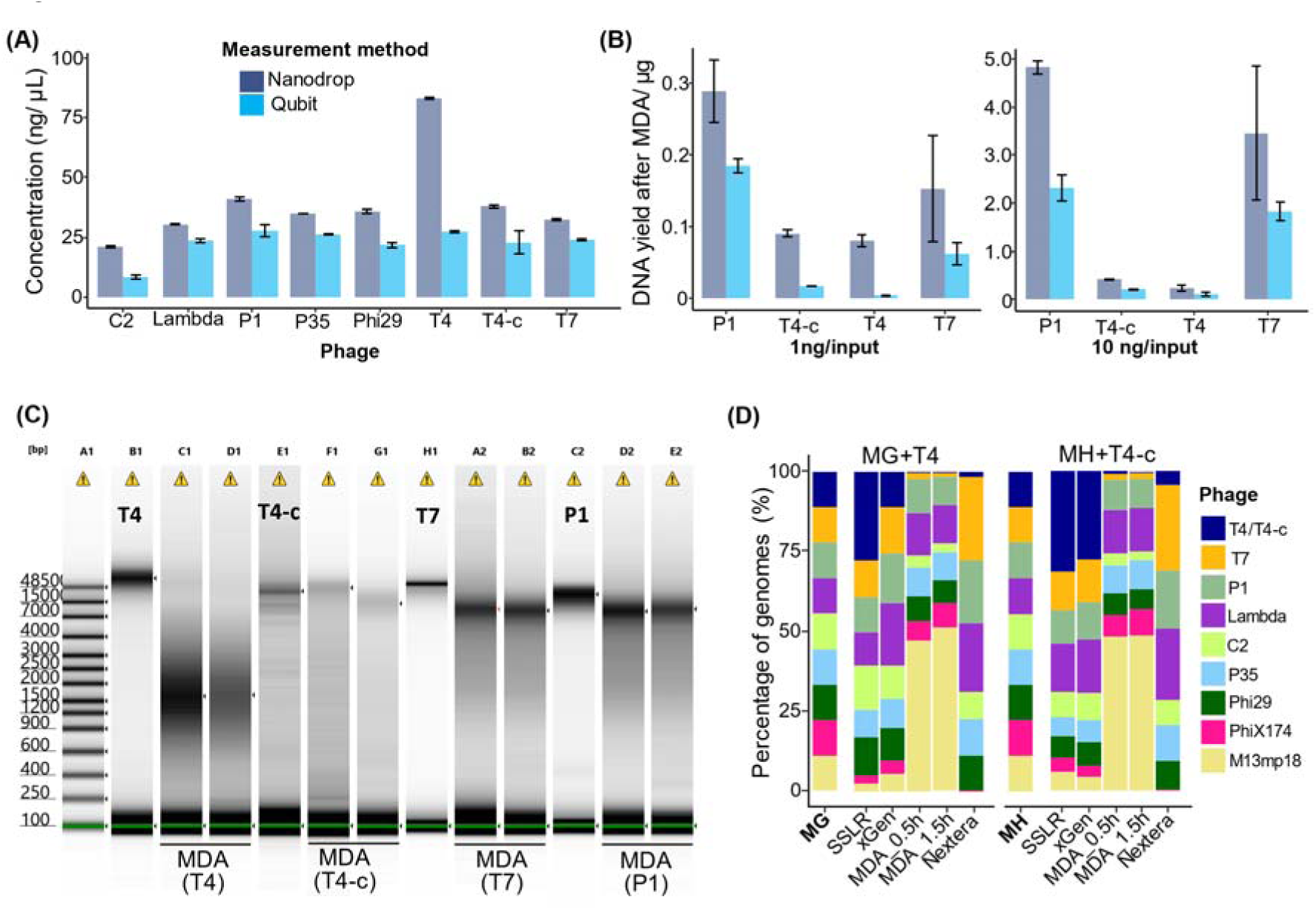
| Quantification and MDA bias for highly modified T4 genome. (A) The concentrations of DNAs used in this study were measured by two different quantification methods (Qubit and Nanodrop). (B) MDA amplification differ in genomes. One and 10 ng of DNA from P1, T4, T4-c and T7 were amplified by 30 min of MDA and purified with Zymol Genomic Purification kit. The y-axis indicates the total yield of amplified MDA products (μg)and measured either by Nanodrop or Qubit. (C) The genomic DNA (T4, T4-c, P1 and T7) and the MDA amplification products (2 repeats for each reaction) were visualized by Tapestation4200 with genomic screentapes. (D) Percentage of phage genomes with T4 or T4-c generated by different library preparation methods. Different color indicate different phage genomes with equally input (11.11%). MG: Mock G contains high modification T4 genome (T4) with equal ratio of all genomes; MH: Mock H contains lower modification T4 genome (T4-c) with equal ratio of all genomes (**Fig. S1G**). The percentage of phage genomes was calculated by aligning the clean reads to customed database (**Supplementary file6**) with bowtie2 to assess the relative abundance of each DNA phage (**Table S5**).

### SSLR can simultaneously identify DNA and RNA viruses

Next, we applied the SSLR method for simultaneous sequencing of DNA and RNA phage genomes (**Table 1**, **Fig. S1B** and **S1F**). We used DMSO or heat treatment to denature the dsRNA to facilitate the reverse transcription (RT) process. We found both treatments including the control (No DMSO) could increase the ratio of ssRNA compared Heat or No Heat, especially for the low ssRNA mock (5% of ssRNA, Mock D) (**Fig. 4A** and **Fig. S7**). Although pretreatment with DMSO increased the number of reads of dsRNA compared to non-DMSO treatment in the mocks with less dsRNA (5% ∼ 25% of dsRNA, Mock D and E, **Table S6**), it was not as effective as heat treatment in terms of read accuracy and sequence coverage (**Fig. 4B**, **Fig. S7** and **Table S6**).

**Fig. 4.**
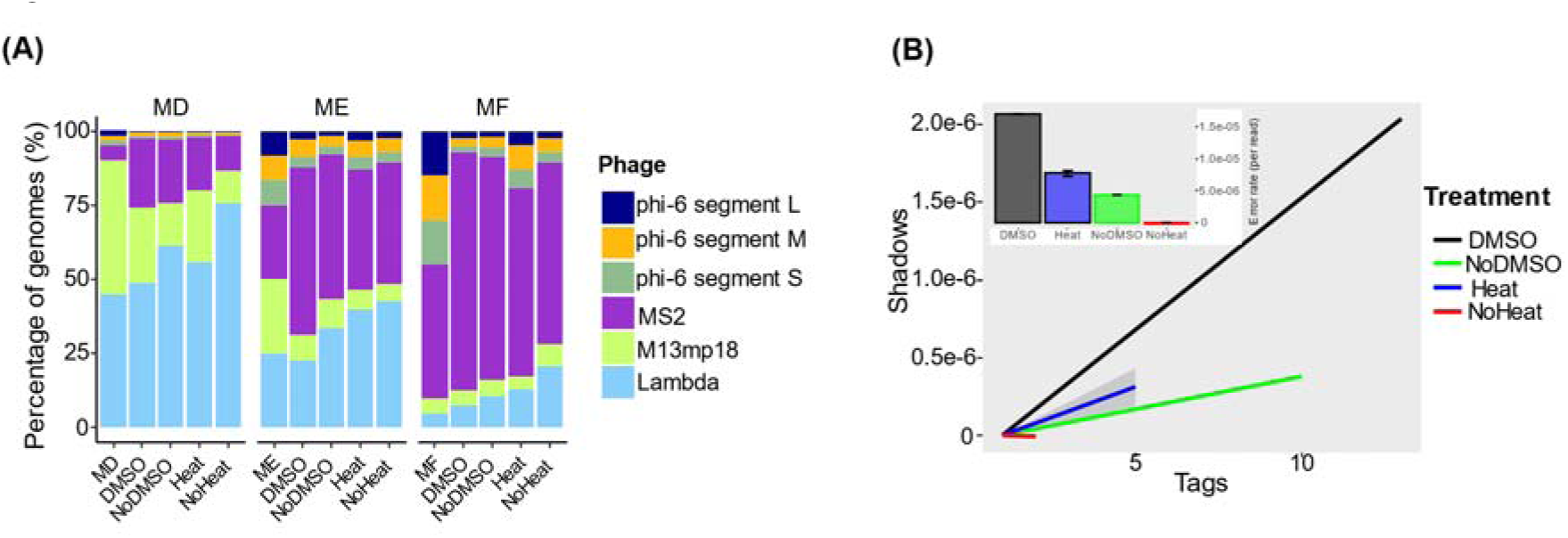
| Comparison of the efficiency of SSLR on the simultaneous identification of DNA/RNA mock communities. SSLR was used to prepare and sequence 4 artificial virome containing different proportions of the DNA phage and RNA phage. (A) Effect of DMSO and heat treatment on the percentage of 4 phage genomes. The percentage of phage genomes was calculated by aligning the clean reads to customed database (**Supplementary file6**) with bowtie2 to assess the relative abundance of each DNA phage (**Table S6**). These phage genome abundance values were calculated based on the quantity of DNA and RNA phages measured by Nanodrop. MD: Mock D with ratio of 90:10 for DNA and RNA; ME: Mock E with ratio of 50:50 for DNA and RNA; MF: Mock F with ratio of 10:90 for DNA and RNA. phi6 has 3 segments, large (6,374 bp), medium (4,063 bp) and small (2,948 bp). (B) Heat treatment does not adversely affect sequencing error rates. The R package ShadowRegression estimates reference-free error rates (inset) based on a transform of the slope of read counts and their ‘shadows’ (main plot line graphs). Shadows (y-axis) are a measure of the variation in read counts across different sequencing runs for the same sample. They are calculated by taking the logarithm of the ratio of read counts in one run to another run. A higher shadow value means a larger difference in read counts between the two runs. Tags (x-axis) are a measure of the abundance of reads for a given nucleotide position in a sample. They are calculated by taking the logarithm of the read count at that position. A higher tag value means a higher number of reads at that position. The figure shows the relationship between shadows and tags for different samples treated with or without Heat/DMSO. The slope of this relationship is used to estimate the sequencing error rate for each sample, which is shown in the inset plots. The figures suggest that DMSO treatment does not affect the sequencing error rate significantly.

**Table 1.**
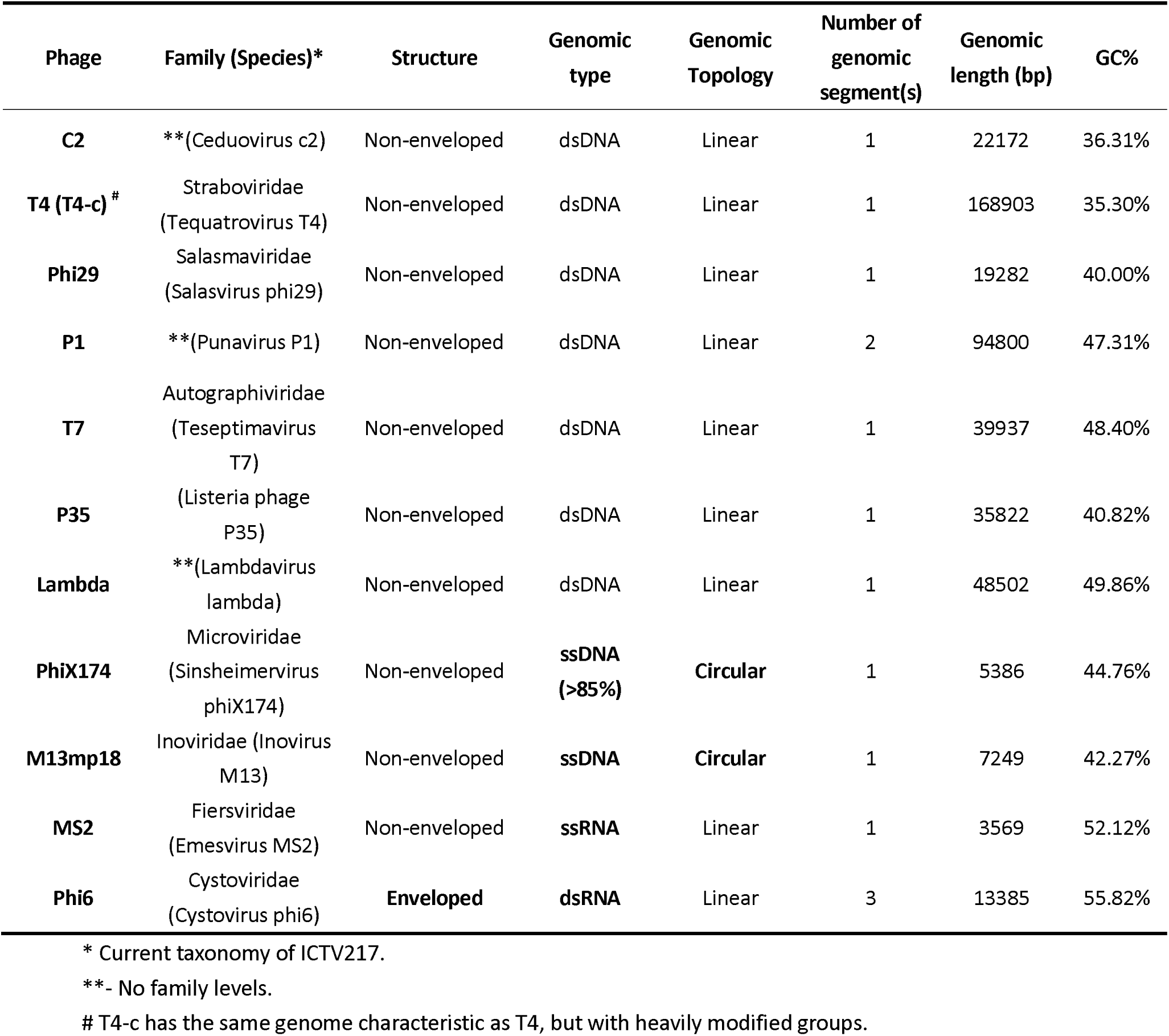
Overview of phage genome characteristics included in the mock communities from the present study.

### SSLR provides high quality fecal virome characterization and allows detection of highly modified phage genome

Finally, we tested the capability for determining fecal virome composition using the methods described above (**Fig. S1D**). Although both MDA based methods yielded more reads that can be assigned to Reference Viral Database (RVDB) (range, 5.77-6.26%, **Fig. S8A** and **Table S7**) compared to SSLR (0.83-1.20%) and xGen (1.23-1.38%) (**Fig. S8B**), 96% of viral operational taxonomic units (vOTUs) were shared among different methods (**Fig. S8C**). We also observed that although the MDA method resulted in higher alignment rates, it led to lower virome Shannon diversity compared to the other library preparation methods (**Fig. S8D** and **8E**). And for both short and long-time MDA, proportions of ssDNA viral families were increased compared to other library preparation methods (**Table S8**). The SSLR could capture around 2-times less of single genome virome (ssDNA of *Microviridae* and ssRNA of *Virgaviridae*) than xGen method (**Table S8**, **Fig. 5B** and **Fig. S8F**).

**Fig. 5.**
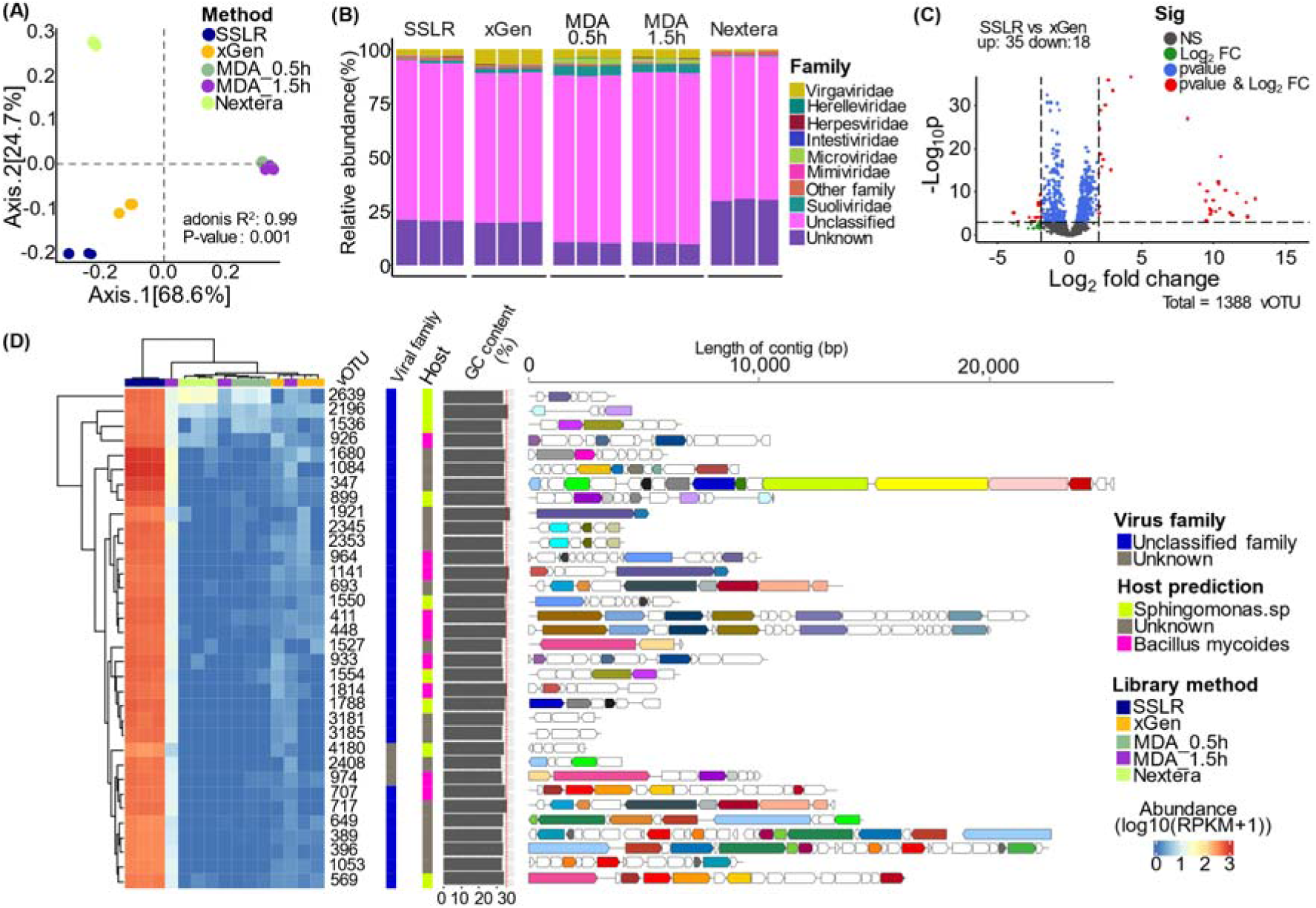
| Comparison of different library strategies for fecal virome communities. (A) Principal coordinates analysis (PCoA) plots of Bray-Curtis distance matrices. PCoA was used to plot the beta diversity of viral-associated communities using the bray matrix. Different colors indicate different library preparation methods. For each axis, in square brackets, the percentage of variation explained was reported. (B) Representative taxonomic distribution (relative abundance) of the sequenced virome. The relative distribution is described at the taxonomical level of the family. The taxonomy of contigs was determined by querying the viral contigs against a database containing taxon signature genes for virus orthologous group hosted at www.vogdb.org. The unclassified are the contigs that cannot be assigned to any known viral taxonomy at the family level, the unknown is the contigs that are related to “viral dark matter”. (C) Volcano plot illustrates differential vOTUs were derived from DESeq2 analysis that had more than or equal to two-fold changes in SSLR versus xGen. Each dot represents a vOTU contig and is colored to indicate significance. Grey: not significant (NS), Green – significant by log2 fold change (>2), Blue: significant by p-value (<0.01), Red – significant by log2 fold change and p-value. Figure made with EnhancedVolcano implemented in R. (D) Heatmap of the high abundance (RPKM) viral contigs that were highly enriched in SSLR method that does not present in the rest of methods. This figure shows a subset of the data presented in **Fig. S8A**. Genomic maps of the open reading frames (ORFs) which are predicted by prodigal and then annotated by the blast from the NCBI protein database, the best hits were used for the annotation. Different colors indicate different annotated protein, directional boxes indicate ORFs in the respective orientation (detailed contigs legend can be found from **Fig. S9D**.

Additionally, the different library preparation methods led to differential abundant core species as compared with the Nextera method although the majority of vOTUs are not identified at family level (**Fig. S8F**). Based on the virome abundance (log10(RPKM+1)), 4 clusters were identified (**Fig. S9A**). Notably, the abundance of cluster 3 from the SSLR method was approximately 10^3^ times higher than the other methods (**Fig. 5D** and **Fig. S9B**). We found that these 34 vOTUs were associated with multiple hits to Sphingomonas phage PAU (previous *Myoviridae*) (**Fig. 5C**, **Fig. 5D** and **Fig. S9C**). As it is evident that the cytosines in Sphingomonas phage PAU genome were highly modified ^38^, this further confirmed that SSLR method can efficiently include modified viral genome in the sequencing libraries.

## Discussion

The viral community in the human gut is highly complex and likely plays an important role in maintaining gut homeostasis ^1, 39^. However, whether the true diversity and distribution of gut virome can be accurately determined by traditional nucleic acids library construction and sequencing methods and with what kind and degree of biases are only partly known. In this study, we addressed these problems by introducing a series of phage genome mock communities reflecting different ratios of four types of viral genomes (dsDNA, ssDNA, dsRNA, ssRNA). To ensure accuracy and avoid extraction efficiency variations ^40, 41^, we used phage genome (DNA/RNA quantified by Nanodrop or Qubit) as initial inputs instead of phage particles by plaque assay or SYBR green counts. The single-stranded library preparation method (SSLR) we introduced for gut virome studies proved to be highly accurate, less biased and comparably efficient, with the added benefits of being fast and cost-efficient.

SSLR demonstrated excellent efficiency in quantifying both dsDNA and ssDNA genomes, overcoming the major limitation of the Nextera protocol, which only works for dsDNA genomes ^7, 42^. While MDA can be applied for turning ssDNA into dsDNA before using Nextera, and it is known for its amplification preference for ssDNA genomes ^23–25^. In line with the previous studies ^24^, we showed that MDA led to 2-3-fold overestimation of ssDNA in low ssDNA mocks (Mock A, around 10% of ssDNA) but not in high ssDNA genome mocks (≥ 50%, Mock B and C). Additionally, the amplification bias appeared to depend on the type of ssDNA genome, as MDA preferentially amplified the alpha3 genome more than PhiX174 and M13mp18, which might be associated to the circular ssDNA of alpha3 being amplified faster than linear ssDNA or the different responses of ssDNA genomes to the heat-denaturing before MDA ^23^. Similar to previous studies we also found that SSLR and xGen slightly underestimated ssDNA in mocks with a low ratio of ssDNA (≤ 10%), this might be due to the ssDNA being more sensitive to shearing during the sonication step compared to dsDNA ^24, 43^.

T4 is an exception among dsDNA genomes in the mocks, as its relative abundance in SSLR and xGen was consistently higher than its input (1.6 – 3.7 fold) and higher than its abundances in Nextera (1.96 – 9.59 fold) and MDA (99 – 586 folds). Interestingly, we observed a significant and consistent difference between the quantification of DNA concentration by Qubit and Nanodrop when measuring T4 genomes. This discrepancy may be attributed to glucosylated 5-hydroxymethyl-cytosine (glu-5HMC) modifications present in the T4 genome ^36, 37^. These modifications could interfere with or alter the binding modes of the Qubit dye, leading to lower concentration values than those obtained through UV absorbance-based quantification (Nanodrop) ^44^. Considering the high quality of our T4 genome, which was extracted, purified, and recovered from agarose gel (see the method part), the concentration values measured from Nanodrop should be more accurate, as 3 times concentration higher from nanodrop is corresponding to the ∼3-times higher sequencing output (**Fig. 3A** vs **Fig. 2A**). Our results also shed light on the reason why the 7-deazaguanine modified Cellulophaga phage phi38:2 (previous *Myoviridae*) was overestimated in the A-LA method (xGen) and underestimated in MDA ^24^, as it was demonstrated recently that this phage has extensive modifications on its genome to protect its DNA from bacterial defense systems ^45^. It was also reported that when measuring potentially highly modified viral DNA using DNA-binding fluorescent dyes, greater caution should be exercised to avoid pitfalls ^46^. In comparison to the T4 genome, the T4-c genome is less modified, and its genome can be sequenced deeper using the Nextera method (**Fig. S6**). However, in the MDA-related method, the genome yield was not increased for T4-c suggesting the remaining modifications in T4-c genome could still inhibit the MDA process on a similar level.

SSLR has an advantage for sequencing highly modified viral genomes (such as T4 with 5-HMC) compared to existing methods, which is further demonstrated by the detection of PAU phage in the fecal virome. We failed to identify PAU-like phage from four gut virome databases (GVD, GPD, MGV and IMG_VR4.1, **Table S10**) ^47–50^, probably due to the modified phages were not included in the original metagenomic sequencing libraries. This finding suggested that many virome studies may have missed or at least underestimated the existence of viruses with highly modified genomes.

Unlike the widely used Nextera library preparation method, which can be affected by genome size and GC-content ^51, 52^, the proposed SSLR protocol exhibits low read errors and provides comprehensive ability for variable genomes with exceptional evenness of coverage and near-complete assembly of phage genomes, as also exemplified for the otherwise “difficult” phage T4. We also observed that SSLR outperforms xGen in several performance metrics (including assembly quality, evenness of genome coverage and error rate, (**Table S4** and **Fig. S3**) probably due to the fact that it does not need synthesizing a second strand for ssDNA viral genomes prior to adaptor ligation, thus retaining the native termini and avoiding possible artifacts or errors presenting during sequencing library preparation ^35^.

SSLR also offers the advantage of using cDNA directly, the first-strand products of RT, as ligation templates for sequencing of RNA viruses. This simplifies the RT step and preserves the strand information ^35^. Our DNA and RNA genome mocks tests demonstrated that the RT step did not significantly affect the distribution of DNA viruses, although the ssRNA virus MS2 was overrepresented. This overrepresentation can be attributed to the faster RT conversion rate of ssRNA compared to dsRNA, leading to non-uniformity in read coverage, which is consistent with previous observations ^53^. Furthermore, for denaturing of dsRNA, the heat-shock treatment appears to be a more cost-effective and easier approach compared to DMSO treatment, it not only reduces the denaturation time but also eliminates the necessity to remove DMSO after treatment, thereby conserving more sample ^53^.

In conclusion, we introduce an improved, rapid, straightforward, and efficient ligation-based single-stranded DNA library preparation method tailored for virome studies.

## Materials and Methods

### Mock communities with phage DNA and RNA genomes

Customized bacteriophage mock community samples were prepared, comprising of a mixture of 4 to 9 bacteriophage genomes with different genome sizes (3.5 to 168.9 kbp), genome types (dsDNA, ssDNA, dsRNA and ssRNA), and G+C content (35 to 56%) (**Table 1**). The phages were propagated using the conditions listed (**Table S10**) and the genomic DNA/RNA extractions were conducted using commercial extraction kits (DNeasy Blood & Tissue Kit for DNA or RNeasy Mini Kit for RNA, both from Qiagen) and purified with GeneJET Genomic DNA Purification Kit (Thermo Scientific, #K0722) for recover the high-quality genome. The quality and integrity of DNA and RNA was checked by agarose gel electrophoresis and Agilent TapeStation 4200 High Sensitivity (HS) RNA ScreenTape. Their concentrations were quantified by dsDNA (or ssDNA) HS Assay Kit or RNA HS assay kit (ThermoFisher Scientific, Waltham, MA, USA) with Qubit4.0, along with Nanodrop1000 (V3.8.1).

### DNA phages library preparation and sequencing

For the DNA phage library preparation, we mixed genomic DNA of 9 phages (7 dsDNAs and 2 ssDNAs) with different ratios (Mock A, Mock B and Mock C) to test 5 different library preparation methods (**Fig. S1A, Fig. S1E** and **Table S1**).

Nextera XT libraries (Nextera) were prepared according to the manufacturer’s protocol (FC-131-1096) (Illumina). Briefly, 1.0 ng of each DNA library was enzymatically fragmented and tagged by tagmentation. Amplification was performed using Illumina dual index (i7 + i5) adapters and cleaned by AMPure XP bead cleanup (A63880l; Beckman Coulter). Two technical replicates were generated for each biological sample, resulting in 6 Nextera-prepared libraries.

The same mixture of phage DNA as mentioned above was amplified by Multiple Displacement Amplification (MDA) using the Genomephi V3 kit (GE Healthcare Life Science, Marlborough, MA, USA) with 10 ng input and incubated at 30°C for 0.5 h (MDA_0.5h) or 1.5 h (MDA_1.5h). Subsequently, the amplified DNA was cleaned using Genomic DNA Clean & Concentrator^TM^ kit (Zymo Research, Irvine, CA, USA). Qubit® 1X dsDNA HS Assay Kit on Qubit Fluorometer (Life Technologies, CA, USA) was used to measure the concentration of the purified DNA. Finally, the library was constructed using the Nextera XT kit (Illumina, San Diego, CA, USA). Two technical replicates were generated for each biological sample, resulting in a total of 12 MDA-prepared libraries.

xGen™ ssDNA & Low-Input DNA Library (xGen, previously known as Swift Bioscience Accel-NGS® 1S) was prepared following the manufacturer’s protocol (Catalog # 10009859, Integrated DNA Technologies, IDT). The Mock (15 ng in 20 µl TE buffer) was randomly fragmented by BioRuptor (Diagenode, Liege, Belgium) at default intensity for 15 sec on and 90 sec off, with 7 cycles at 4°C as described by Hoang et al ^55^. Sheared DNAs were firstly subjected to 3’ end tailing and ligation with adapter1, followed by extension and ligation with adapter2. The purified ligation products were indexed with xGen™ Unique Dual Index (UDI) Primers (Catalog # 10005975, IDT) and amplified through a 10-cycle PCR. The amplified products were cleaned up with AMPure XP beads (Beckman Coulter Genomic, CA, USA). For each biological sample, two technical replicates were conducted, resulting in a total of 6 xGen-prepared libraries.

For the Single-stranded library preparation (SSLR, detailed in **Supplementary file1**), the same mocks were used. After random fragmentation, the end modified adapters (**Supplementary file2**) synthesized by IDT (IDT, Leuven, Belgium) were ligated to DNA fragments employing the method from Troll et al ^35^. Subsequently, the ligation mix was purified using MinElute Reaction Kit (Qiagen, Hilden, Germany) and the ligation products were eluted with 23 µl of TE buffer (10 mM, pH8.0). The cleaned ligation products were then amplified and indexed for sequencing with Illumina dual Indexes (i7 + i5) and further purified using AMPure XP beads (Beckman Coulter Genomic, CA, USA). An additional bead cleanup step can be applied to mitigate the formation of byproducts and adapter dimers resulting from self-ligation of the adapters (**Fig. S10**). Two technical replicates were performed for each biological sample, resulting in a total of 6 SSLR-prepared libraries. A detailed protocol can be found on (**Supplementary file1**).

The DNA concentrations of Nextera-prepared, MDA-prepared, xGen-prepared and SSLR-prepared libraries were measured by Qubit® 1X dsDNA HS Assay Kit on Qubit Fluorometer (Life Technologies, CA, USA) and fragment length distributions were determined using TapeStation 4200 (Agilent, CA, USA). Equal amounts of DNA from each library (except xGen-prepared) were pooled and sequenced using 2×150 bp paired-end settings on an Illumina NextSeq550 platform (Illumina, CA, USA). The xGen-prepared libraries were sequenced separately on an iSeq100 System (Illumina, CA, USA) using 2×150 bp paired-end settings due to their indexes being incompatible with the Illumina index.

### RNA phages library preparation and sequencing

Four types of phage genome (**Table 1**, dsDNA, ssDNA, dsRNA and ssRNA) with different ratios were prepared for phage DNA/RNA mock communities (Mock D, Mock E and Mock F). We used three mock communities to determine the applicability of the SSLR method for quantification of RNA and DNA phages simultaneously (**Table 1, Fig. S1B** and **Fig. S1F**). Specifically, 20 µl mock community was treated with DMSO at a final concentration of 50% and incubated at 65 °C for 90 min ^53^. The DMSO was subsequently removed using a QIAmp viral RNA mini kit (Qiagen, Hilden, Germany), according to the manufacturer’s instructions. As a control, a DMSO-free treatment was included. Alternatively, the same mock community was denatured at 95°C for 3 min and immediately snap-cooled on ice, maintaining the nucleic acids as single-stranded. Again, a sample without heat treatment was included as a control library.

All the DMSO-treated samples, heat-treated samples and their corresponding controls were subjected to reverse transcription in a 20-µl reaction system according to the user guide (SuperScript™ IV VILO™ Master Mix, Invitrogen™) and the reaction mix was purified using DNeasy Blood & Tissue Kits (Qiagen, Hilden, Germany) after RT with modifications, where the purification procedure started from step 3, whereby an equal volume of ethanol (96%) was added to the PCR reaction, and a 20 µl of Tris buffer (10 mM, pH8.0) was used for the final elution. Finally, the library preparations with SSLR were performed according to the above-mentioned steps and sequenced on a NovaSeq6000 platform (Illumina, CA, USA).

### Sequencing of highly modified phages

Two types of DNA phage mock communities (Mock G and Mock H) were prepared with equal input (11.11% for each of the 9 genomes, **Fig. S1C** and **Fig. S1G**). Mock G contained a highly modified T4 genome (T4), while Mock H contained a less modified T4-c genome (T4-c). All the libraries were prepared as described above and sequenced with the NovaSeq6000 platform (Illumina, CA, USA).

### Human fecal virome isolation and sequencing library preparation

For environmental sample, a fresh fecal sample was obtained from an anonymous healthy adult (Ethical Committee of the Capital Region of Denmark registration number H-20028549) and thoroughly mixed with SM buffer. Fecal virome isolation and purification were carried out according to our previous method with the following modifications ^16^: the Centriprep 50K was replaced by Centrisart® I centrifugal ultrafiltration unit (MWCO 100 kDa, Sartorius Stedim Biotech GmbH) and the enrichment step was conducted at 2500 × g for 30 min at 4°C. The QIAmp viral RNA mini kit (Qiagen, Hilden, Germany) was used for the extraction of viral DNA/RNA from the concentrated virome solution (**Fig. S1D**). The library preparations were conducted with the aforementioned methods in triplicate. An equal amount of DNA from each library was pooled and sequenced using 150 bp paired-end settings on an NextSeq550 platform (Illumina, CA, USA).

### Metavirome sequencing, data pre-processing and data analyses

Metavirome reads were quality filtered, trimmed and assembled using a previously published pipeline from GitHub: https://github.com/jcame/virome_analysis⍰FOOD. Detailed data analyses for sequencing of the mock communities and fecal virome can be found from **Supplementary file3**. For mock community data, plots were generated using the ggplot2 ^56^ package in R (v4.2.0). Analyses of viral community α- and β-diversity were performed using packages Phyloseq (v1. 36.0) ^57^ and Vegan (v2.6.2) ^58^ in R. For α-diversity analyses, indexes of observed taxa and Shannon diversity were calculated with t.test using the package ggsignif (v0.6.3) ^59^. For β-diversity analyses, Bray-Curtis distance metrics were calculated and unconstrained ordination was performed using principal coordinate analysis (PCoA). To identify differentially enriched vira on the summarized family level, DESeq2 (v1.42.0) was adopted ^60^. The results were then visualized in a heatmap using the R package complexheatmap (v2.18.0) ^61^. Estimation of sequencing error rates was conducted according to Wilcox et al ^53^. Coverage biases were visualized using the python script from: https://github.com/padbr/gcbias, with modifications made by adding the Pearson correlation. To visualize the functional genes of annotated viral contigs, DnaFeaturesViewer (v3.1.0) ^62^ and R package gggenomes (v 0.9.12.9000) ^63^ was used.

## Data availability

The authors declare that the data supporting the findings of this study are available within the paper and its supplemental information files. All the raw viral metagenome sequences data produced in this study are available through the National Center for Biotechnology Information’s Sequence Read Archive under BioProject accession number PRJNA1094595 with submission No. SUB13375064 (DNA mocks), No. SUB14350434 (DNA/RNA mock), No. SUB14350532 (Modification mock), and No. SUB14350824 (fecal virome).

The public databases for checkV (v1.5) at https://portal.nersc.gov/CheckV, VIBRANT (v1.2.1) at (https://github.com/AnantharamanLab/VIBRANT/tree/master/databases), Virsoter2 at (https://osf.io/u3t4j), VOG217 at (https://fileshare.csb.univie.ac.at/vog/vog217) and Iphop (v1.3.2) at (https://portal.nersc.gov/cfs/m342/iphop/db/) are accessible online.

## Code availability

This paper does not generate the original R code for data analysis and visualization.

Code for metavirome pre-analysis is available from GitHub: https://github.com/jcame/virome_analysis-FOOD.

Original python code from (https://github.com/padbr/gcbias) with modifications for fitting the GC bias can be found from GitHub: https://github.com/padbr/gcbias.

The read accuracy and mutation analysis can be found on GitHub: https://github.com/awilcox83/dsRNA-sequencing.

Further information and requests for data, code and resources required to reanalyze the data reported in this paper should be directed to and will be fulfilled by XC. Zhai and L. Deng.

## Author information

Authors and Affiliations

**Section for Food Microbiology, Gut Health and Fermentation, Department of Food Science, University of Copenhagen, Rolighedsvej 26, 1958 Frederiksberg C, Denmark**

Xichuan Zhai, Lukasz Krych, Dennis Sandris Nielsen, Ling Deng

**Section of Microbial Ecology and Biotechnology, Department of Plant and Environmental Sciences, University of Copenhagen, Thorvaldsensvej 40, 1871 Frederiksberg C, Denmark**

Alex Gobbi, Witold Kot

**European Food Safety Agency (EFSA), Via Carlo Magno 1A, 43126 Parma, Italy**

Alex Gobbi

## Contributions

This research was designed and directed by LK, DSN and LD. The experiment was performed by XZ. Data collection was performed by XZ, AG and WK. Data analysis was done by XZ and LD. The manuscript was written and revised by XZ, DSN and LD. All authors read and approved the final manuscript.

## Corresponding authors

Correspondence to Ling Deng

## Additional data

**Additional file 1**: SSLR adapter design.

**Additional file 2**: Step-by-step library preparation protocol for the reproposed SSLR.

**Additional file 3**: Metavirome sequencing and data pre-processing.

**Additional file 4**: Supplementary Table S1-S10. **Table S1**: Summary of DNA genome sequences. Top table: Relative abundance and coverage of phage genomes in libraries from DNA mock communities (Mock A, Mock B and Mock C) with T4 genome. The relative abundance of each genome was calculated based on its coverage by using bowtie2 alignment with our customed database (**Additional file6)**. For comparing the efficiency of different methods for each group of phages (i.e., dsDNA and ssDNA), relative abundance of two type genomes were computed from this table (by dividing the value for each phage by the sum for the group). Middle table: The average coverage for each genome from each library preparation method. Bottom table: similar to top table without T4 genome. **Table S2**: Comparisons of cost and effectiveness of Mock virome library preparation method. This comparison only considers the library preparation step for DNA virome. **Table S3**: Overview of metavirome sequencing for the DNA phage mock community. Statistics from assembly before and after quality checks with CheckV, vibrant and virsorter2. Contigs from each library prepared methods were evaluated by quality checks with CheckV, vibrant and virsorter2 and then subjected to the Quast for quality assessment of assembled contigs. The theatrical statistics of mock community were also subjected to Quast and bolded at the bottom of table. **Table S4**: Summary of contigs hits to the customed mock database. The quality-checked contigs were subjected to the customed database and the hit number was counted and listed in the table. **Table S5**: Summary of DNA genome sequences with interested modified genomes (T4 and T4-c). T4 is a highly modified genome, T4-c has lower modification compared to T4 genome. **Table S6**: Summary of DNA/RNA genome sequences prepared with SSLR method. Relative abundance and coverage of phage genomes in libraries from DNA/RNA mock communities (Mock D, Mock E and Mock F) with different treatments (heat, no heat, DMSO and no DMSO). The relative abundance of each genome was calculated based on its coverage by using bowtie2 alignment with our customed database. For comparing the efficiency of different treatments for each type of phage genome (DNA and RNA), relative abundance of two type genomes (DNA or RNA) were calculated from this table (by dividing the value for each phage by the sum for the genome types). Middle table: The average coverage for each genome from each library preparation method. The average coverage for each genome from each treatment method is listed in the bottom table. **Table S7**: Taxonomic classification of fecal metavirome sequencing reads into the categories viral, human, bacterial, and unknown origin. To check the presence of non-viral DNA sequences, 50,000 random forward reads were used according to their match to a range of viral, bacterial, and human reference database of Kaiju 1.8.2. **Table S8**: Relative abundance of fecal virome prepared with 5 different library strategies at taxonomy of family level (support data for **Fig. 5B**) and the overall alignment rate of different library preparation methods based on bowtie2 alignment to the vog217 database. **Table S9**: Top 5 hits of PAU phage on 4 virus databases (GVD, GPD, MGV and IMG_VR4.1). **Table S10**: Characteristics of phage genomes included in the mock communities from the present study, as well as growth conditions for the strains in the mock communities. **Table S11**: Average alignment rate of each library to the customed databases for all the mock communities tested in the present study.

**Additional file 5**: Supplementary figures S1-S10.

## Supporting information

Supplemental file1_Virome library workflow

Supplemental file2_Adapters design

Supplemental file3_Data analysis

## Acknowledgements

We thank the donors and our colleagues at Section for Food Microbiology, Gut Health and Fermentation, Department of Food Science, University of Copenhagen (KU FOOD) for their participation and cooperation. We acknowledge Dr. Samuel Kilcher and Prof. Martin Loessner from Institute of Food, Nutrition, and Health, ETH Zurich for providing phage P35 and its host strain *Listeria monocytogenes* Mack. We acknowledge Dr. Yuvaraj Bhoobalan-Chitty from Department of Biology, University of Copenhagen (Ole Maaløes Vej 5, 2200 Copenhagen N, Denmark) for providing phage T4-c. XZ was supported by China Scholarship Council (CSC) Grant #201906870027.

## Abbreviations

SSLR: single-stranded library; Single Reaction Single-stranded LibrarY: SRSLY; RT: Reverse transcription; MDA: multiple Displacement Amplification; PCoA: principal coordinates analysis; vOTUs: viral-operational taxonomic units; ORFs: open reading frames; NCLDV: nucleocytoplasmic large DNA viruses; LCA: Lowest Common Ancestor; SGBs: species-level genome bins; NS: not significant; gut virome database: GVD; gut phage database: GPD; metagenomic gut virus: MGV; high-Integrated Microbial Genomes /Viruses 4.1: IMG/VR4.1; High Sensitivity: HS; Unique Dual Index: UDI; Integrated DNA Technologies: IDT

## Ethics approval and consent to participate

A healthy adult anonymous donor donated fecal samples for the study after being informed orally and in writing and provided written consent (Ethical Committee of the Capital Region of Denmark registration number H-20028549).

## Consent for publication

Not applicable.

