## Supplemental Figures for "A single strand-based library preparation method for unbiased virome characterization"

Supplementary figures for the SSLR paper entitled " A single strand-based library preparation method for unbiased virome characterization"

By

Xichuan Zhai, Alex Gobbi, Witold Kot, Lukasz Krych, Dennis Sandris Nielsen, Ling Deng

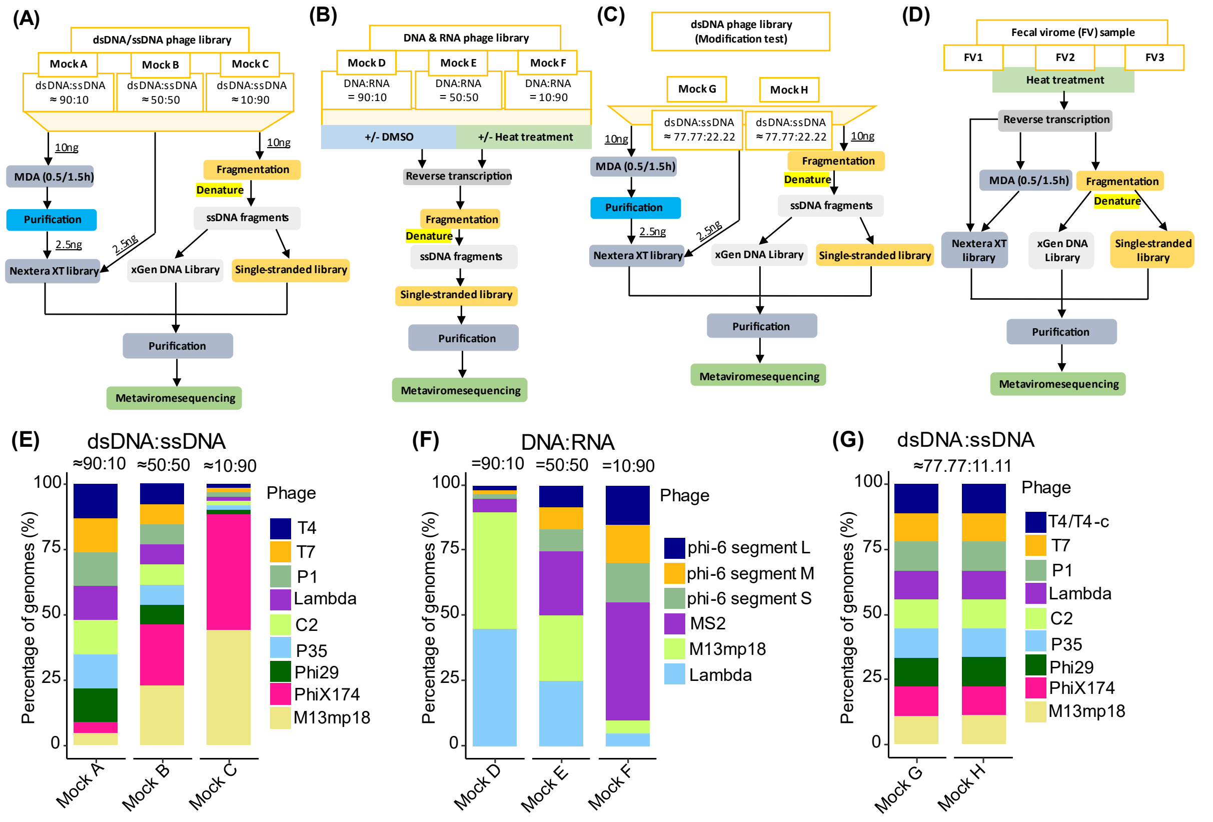


**Fig. S1** | Virome sequencing library preparation strategy with different types of nucleic acids (dsDNA, ssDNA, dsRNA and ssRNA). (A) Workflow for DNA phage genomes (dsDNA and ssDNA) library preparation with different library methods to varying ratios of genomes (Mock A, Mock B and Mock C). (B) Workflow for DNA and RNA phage genomes library preparation with the single-stranded library at different ratios of genomes (Mock D, Mock E and Mock F). (C) Workflow for comparison of the modification existence for (Mock G with T4, Mock H with T4-c), where T4 is a heavily modified genome and T4-c is a lower modified genome. (D) Workflow for comparison of the fecal sample (Fecal virome: FV) with different library preparations. (E) Percentage of phage genomes of DNAs library with different ratios of ssDNA and dsDNA phage genomes, three artificial virome communities (Mock A, Mock B and Mock C) containing different proportions of the ssDNA phage PhiX174 and M13mp18 mixed with dsDNA phage. These phage genome abundance values were calculated based on the quantity of dsDNA and ssDNA phages measured Qubit dsDNA (or ssDNA) HS Assay kit. Mock A: ratio of ~90:10 for dsDNA and ssDNA; Mock B: ratio of ~50:50 for dsDNA and ssDNA; Mock C: ratio of ~50:50 for dsDNA and ssDNA. (F) Percentage of phage genomes of DNA/RNAs library with different ratios of DNA and RNA phage genomes, three artificial viromes containing different proportions of the dsRNA phage Phi6 and ssRNA MS2 mixed with DNA phage. These phage genome abundance values were calculated based on the quantity of DNA and RNA phages measured with Nanodrop1000. Mock D: ratio of 90:10 for DNA and RNA; Mock E: ratio of 50:50 for DNA and RNA; Mock F: ratio of 10:90 for DNA and RNA. (G) Percentage of phage genomes of DNAs library with 2 different T4 genomes, where MG: Mock G containing high modification T4 genome (T4) with equal ratio of all genomes; MH: Mock H containing lower modification T4 genome (T4-c) with equal ratio of all genomes.

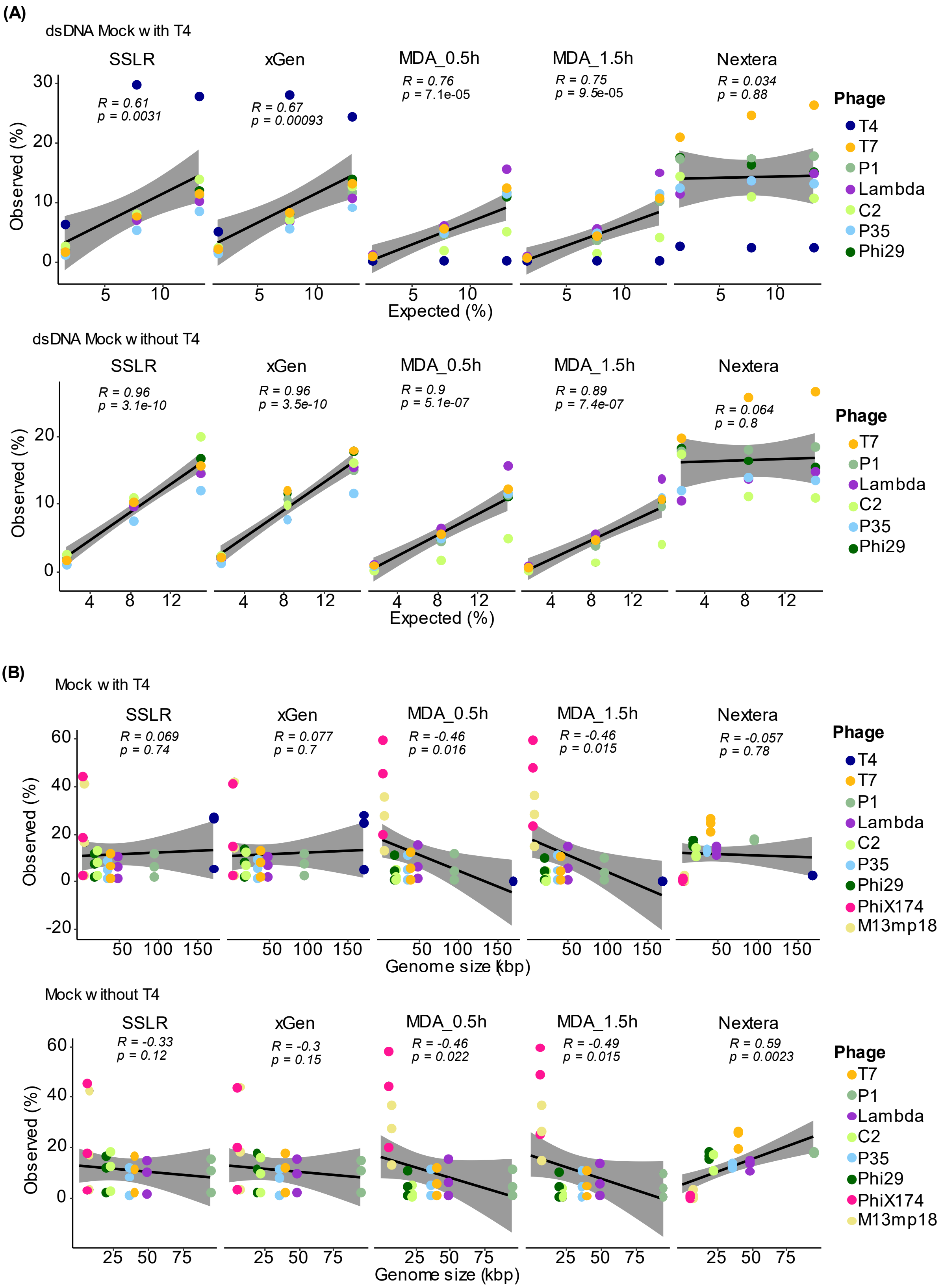
(A)
dsDNA Mock with T4
SSLR
xGen
MDA_0.5h
MDA_1.5h
Nextera
R = 0.61
p = 0.0031
R = 0.67
p = 0.00093
R = 0.76
p = 7.1e-05
R = 0.75
p = 9.5e-05
R = 0.034
p = 0.88
Observed (%)
Expected (%)
Phage
T4
T7
P1
Lambda
C2
P35
Phi29
dsDNA Mock without T4
R = 0.96
p = 3.1e-10
R = 0.96
p = 3.5e-10
R = 0.9
p = 5.1e-07
R = 0.89
p = 7.4e-07
R = 0.064
p = 0.8
(B)
Mock with T4
R = 0.069
p = 0.74
R = 0.077
p = 0.7
R = -0.46
p = 0.016
R = -0.46
p = 0.015
R = -0.057
p = 0.78
Genome size (kbp)
PhiX174
M13mp18
Mock without T4
R = -0.33
p = 0.12
R = -0.3
p = 0.15
R = -0.46
p = 0.022
R = -0.49
p = 0.015
R = 0.59
p = 0.0023

(C)

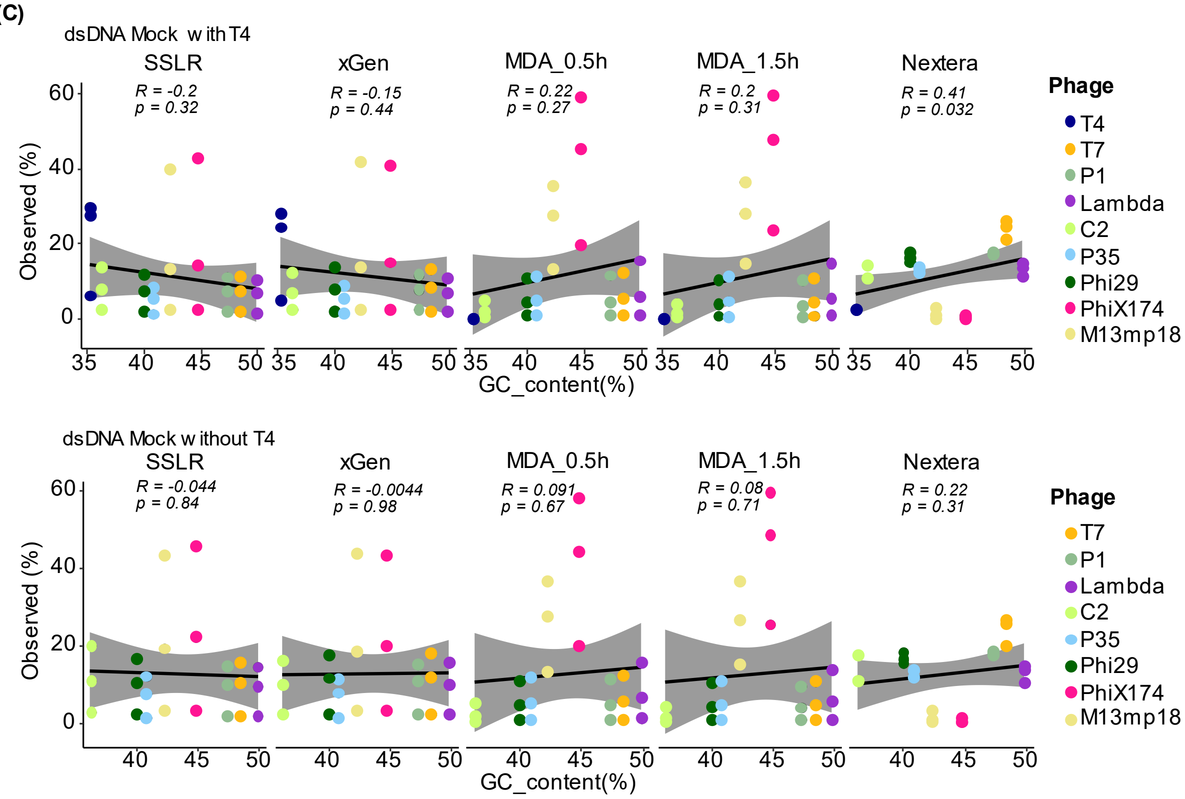


**Fig. S2** | Pearson correlation coefficient (r) and two-tailed p-value between the expected and obtained read distributions (in percentage) in different DNAs phage mock communities with or without phage T4 genome. (A) Pearson correlation only dsDNA phage genome with (Top panel) or without T4 (bottom panel) shows that the presence of T4 distorted the percentage of dsDNA phage genomes. The phage genome size (B) and GC_content (C) have minimal effect on the phage genome distributions for the library prepared with SSLR and xGen.

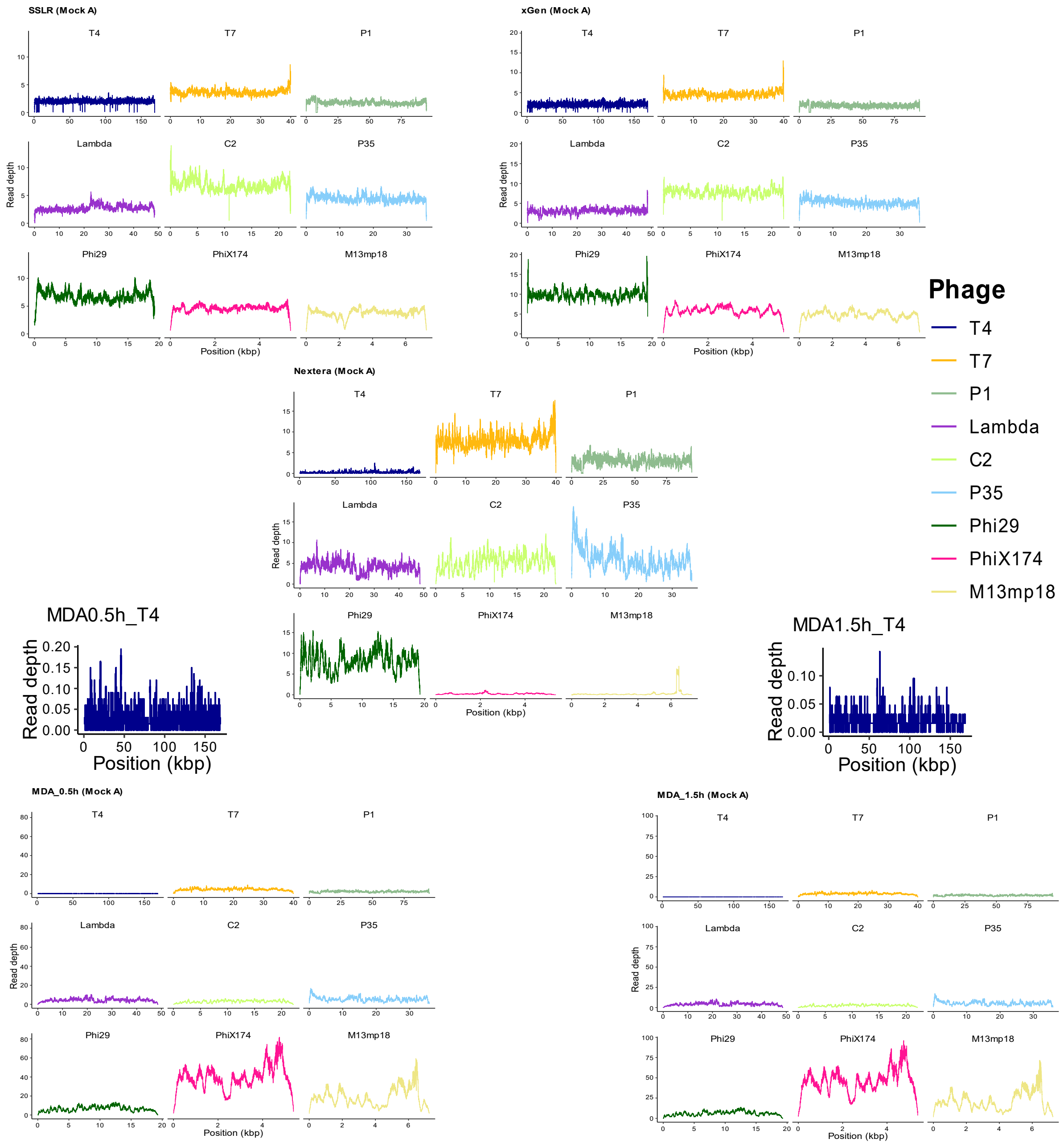


**Fig. S3** | Different library methods induce different sequence coverages of the DNA phage genomes. Read depth at each position in the phage genome was normalized by count/total reads*10000 for easier comparison among different library preparation methods.

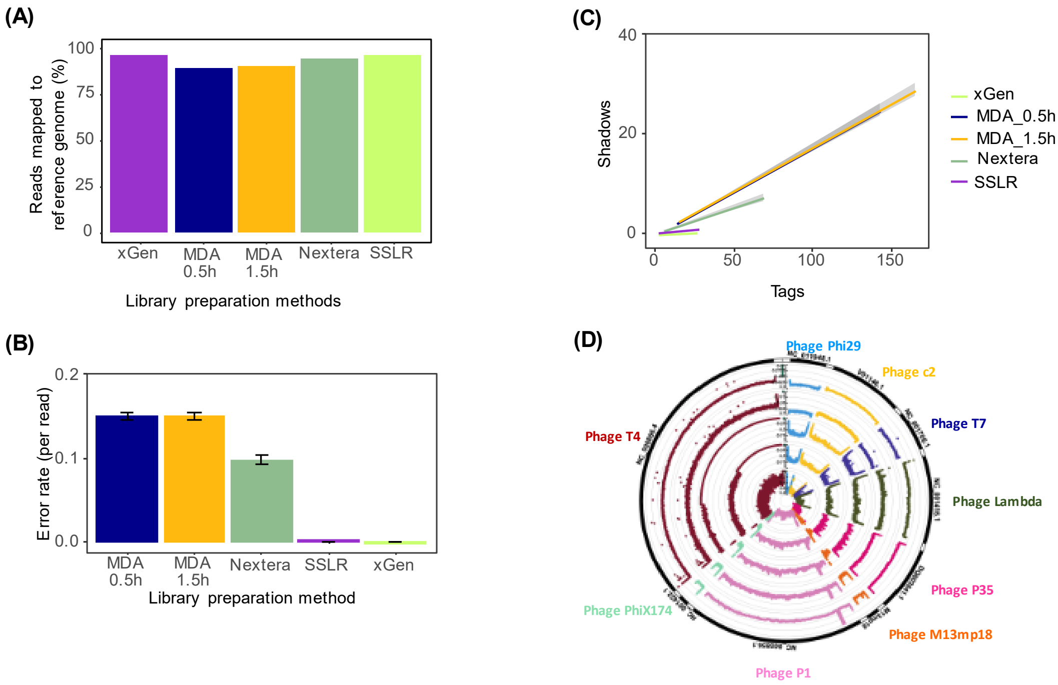


**Fig. S4 |** (A) The overall percentages of sequencing reads mapping to the 9 DNA reference genomes. The R package ShadowRegression estimates reference-free error rates (B) based on a transform of the slope of read counts and their 'shadows' (C). (D) Depth biases in the sequencing of DNA phage genomes prepared with different library methodsThe circle plot shows from the inside: Nextera (Ring 1); MDA_0.5h (Ring 2); MDA_1.5h (Ring 3); xGen (Ring 4) and SSLR (Ring 5). The circles are numbered from the inside. The sequencing depth was shown in the log10 scale. All the reads were nomalized and then analyzed the .

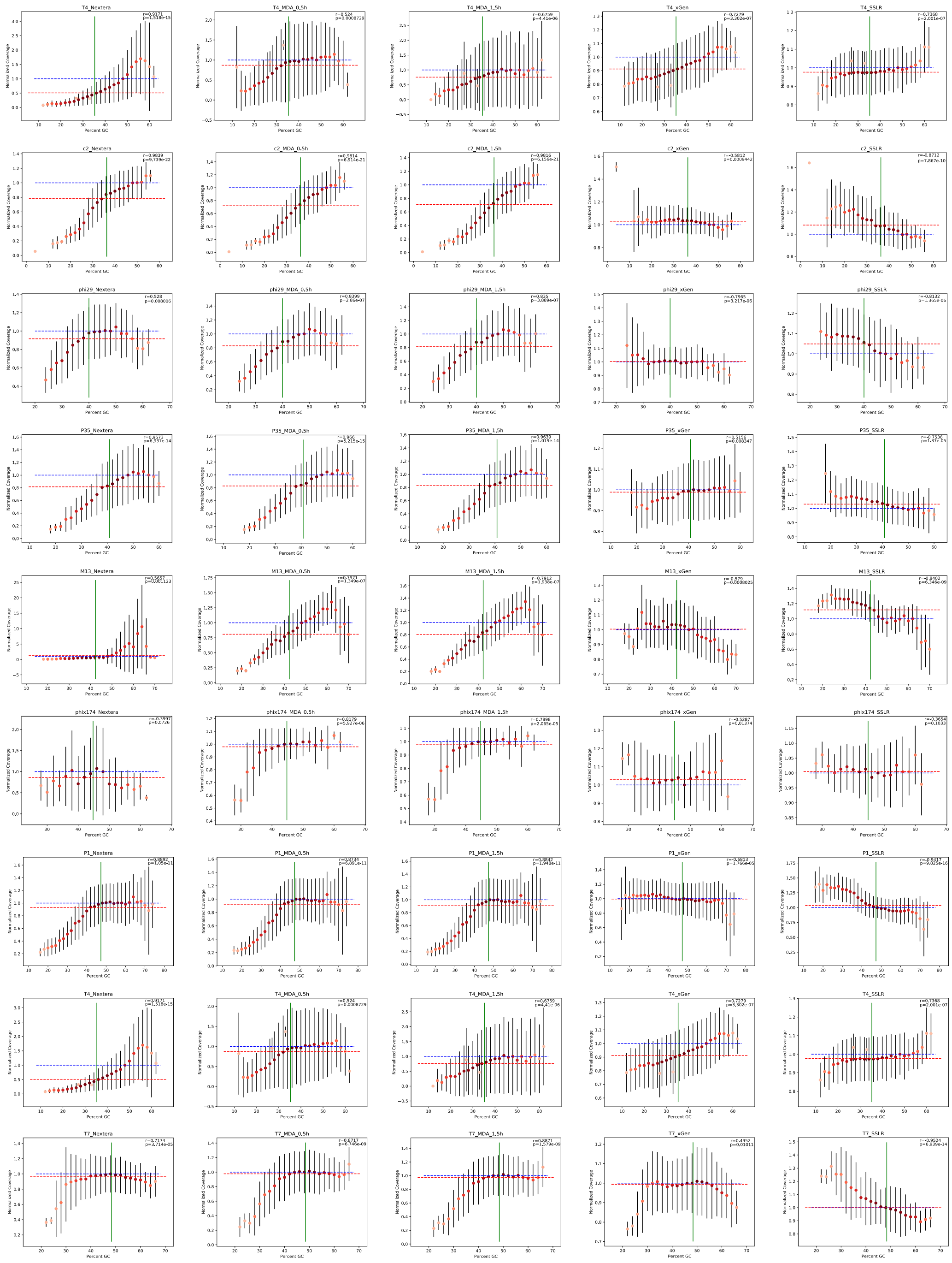

**Fig. S5** | Coverage biases in datasets for the different DNA phage genomes with different GC contents. Dot plots show local GC content and normalized relative coverages in 50-nt windows of NextSeq/iSeq and data from a variety of phages with different average GC contents. Error bars indicate ±1 standard deviation of normalized coverage. The intensity of the blue in the dots is a log-transformed heat map of the number of 50-nt windows averaged into that datapoint. The datapoint with the most windows in each plot has maximum red. The vertical green line marks the average GC content of each assembly. The average normalized coverage value is indicated with a horizontal dashed red line. The plots were visualized with the python script from: https://github.com/padbr/gcbias, with the modifications by adding the Pearson correlation coefficient (r), two-tailed p-value (top right corner of each plot).

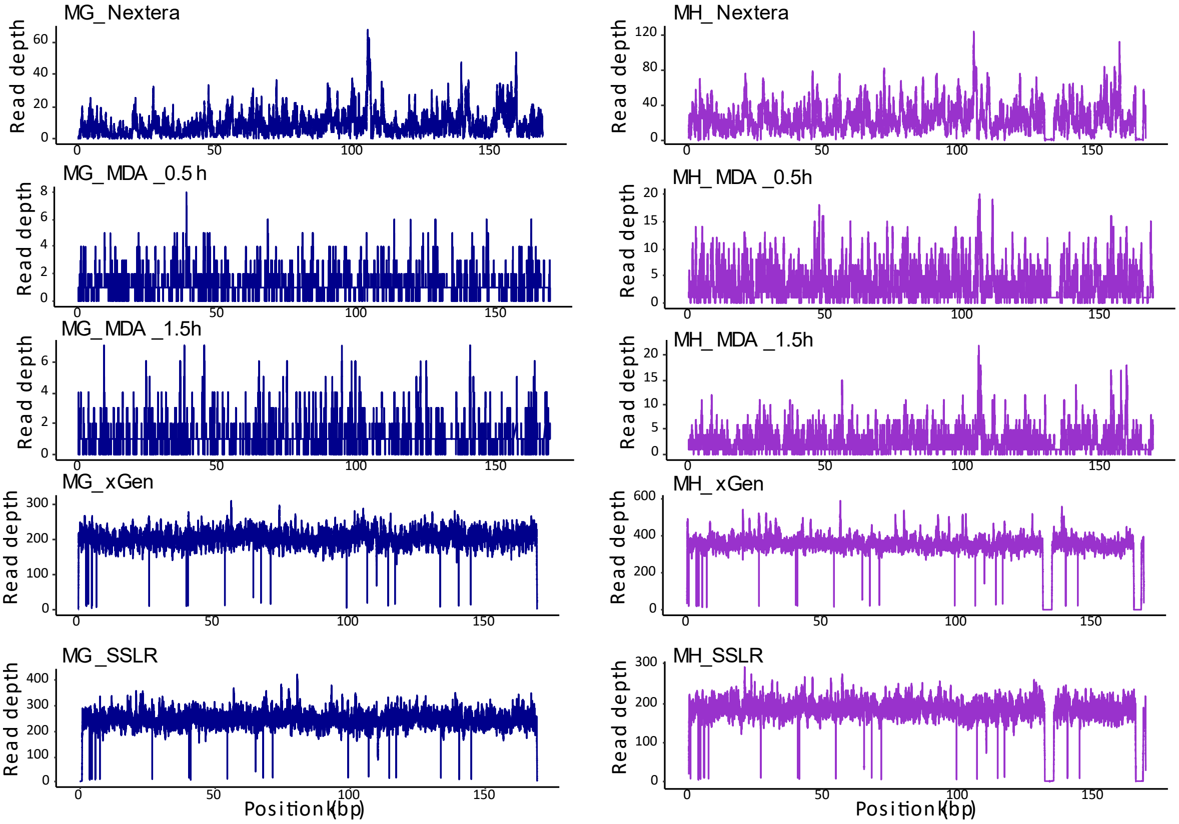


**Fig. S6** | Different library methods induce different sequence coverages of the T4 (blue line)/T4-c (purple line) phage genomes. Read depth at each position in the phage genome was normalized by count/total reads*10000 for easier comparison among different library preparation methods.

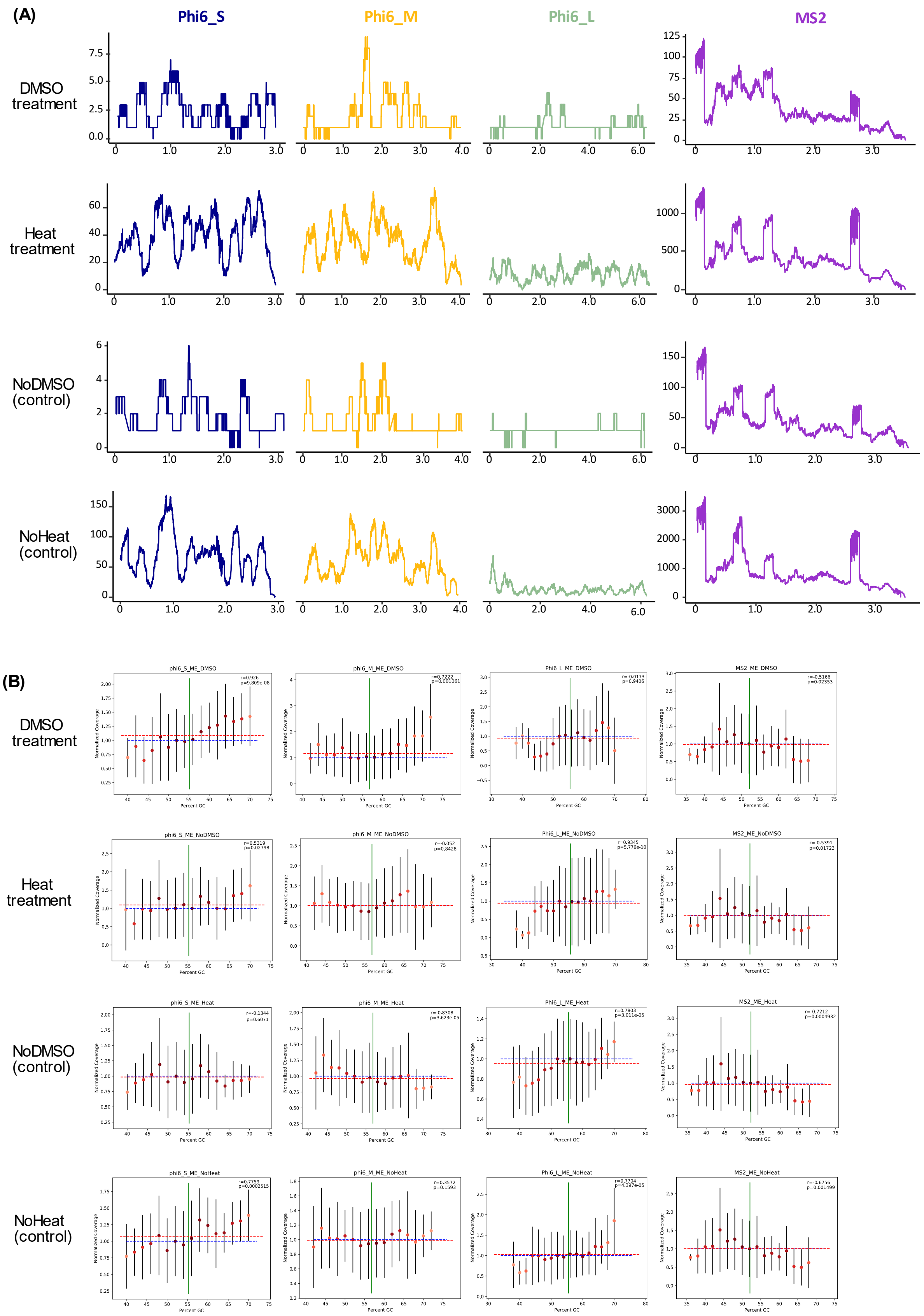

(A)
Phi6_S
Phi6_M
Phi6_L
MS2
DMSO treatment
Heat treatment
NoDMSO (control)
NoHeat (control)
(B)
DMSO treatment
Heat treatment
NoDMSO (control)
NoHeat (control)
phi6_S_ME_DMSO
phi6_M_ME_DMSO
Phi6_L_ME_DMSO
MS2_ME_DMSO
phi6_S_ME_NoDMSO
phi6_M_ME_NoDMSO
Phi6_L_ME_NoDMSO
MS2_ME_NoDMSO
phi6_S_ME_Heat
phi6_M_ME_Heat
Phi6_L_ME_Heat
MS2_ME_Heat
phi6_S_ME_NoHeat
phi6_M_ME_NoHeat
Phi6_L_ME_NoHeat
MS2_ME_NoHeat
Normalized Coverage
Percent GC

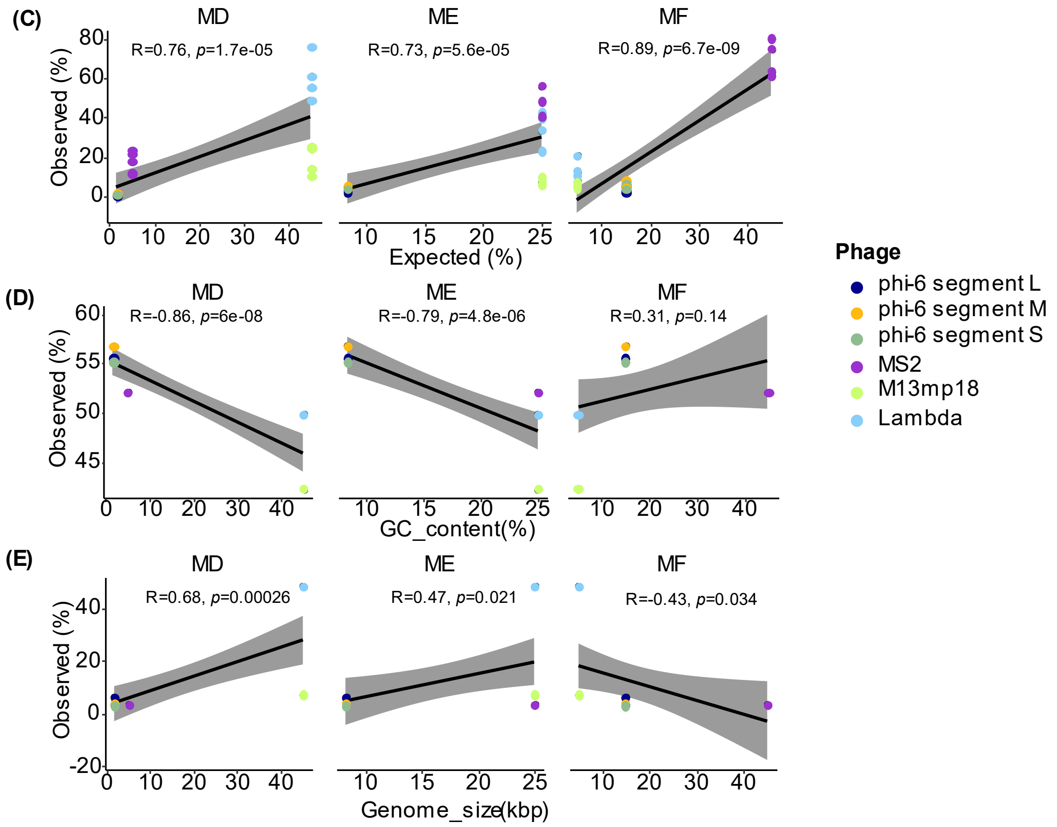


**Fig. S7** | (A) Effect of DMSO and heat treatment on sequence coverage of the RNA phage genomes. Read depth at each position in the phage genomes under varying different treatments (DMSO and Heat) and their controls (NoDMSO and NoHeat) are shown. (B) Coverage biases in datasets for the different types of RNA phage genomes with different GC contents. Dot plots show local GC content and normalized relative coverages in 50-nt windows and data from a variety of phages with different average GC contents. Error bars indicate ±1 standard deviation of normalized coverage. The intensity of the blue in the dots is a log-transformed heat map of the number of 50-nt windows averaged into that datapoint. The datapoint with the most windows in each plot has maximum red. The vertical green line marks the average GC content of each assembly. The average normalized coverage value is indicated with a horizontal dashed red line. The plots were visualized with the python script from: https://github.com/padbr/gcbias, with the modifications by adding the Pearson correlation coefficient (r), two-tailed p-value (top right corner of each plot). (C) Pearson correlation coefficient (r) and two-tailed p-value between the expected and obtained read distributions (in percentage) in different DNA/RNA phage mock communities. The phage genome size (D) and GC_content (E) have minimal effect on the phage genome distributions for the library prepared with SSLR.

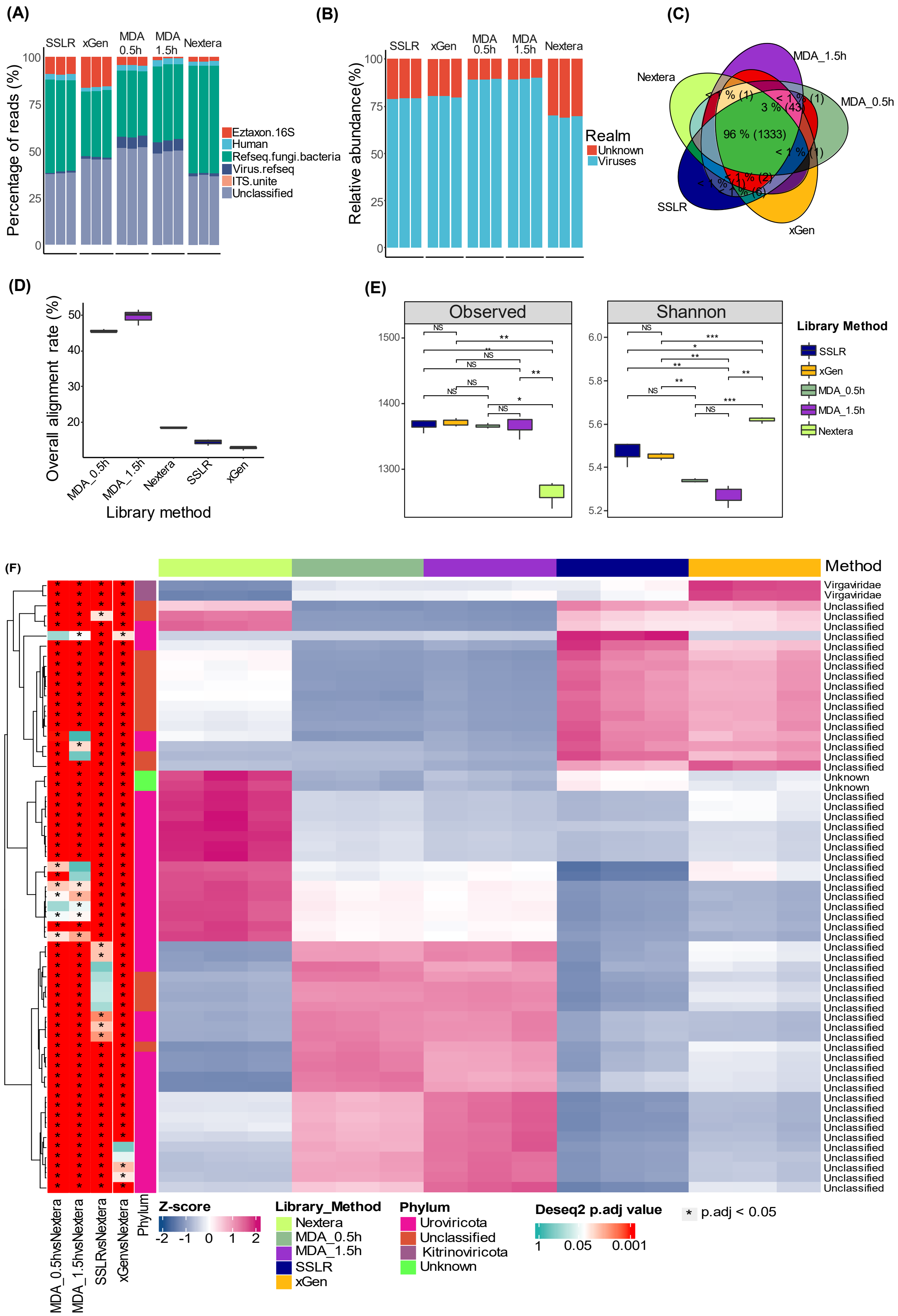
(A)
SSLR
xGen
MDA 0.5h
MDA 1.5h
Nextera
Percentage of reads (%)
Eztaxon.16S
Human
Refseq.fungi.bacteria
Virus.refseq
ITS.unite
Unclassified
(B)
Relative abundance(%)
Realm
Unknown
Viruses
(C)
MDA_1.5h
Nextera
MDA_0.5h
SSLR
xGen
96 % (1333)
3 % (43)
(D)
Overall alignment rate (%)
Library method
MDA_0.5h
MDA_1.5h
Nextera
SSLR
xGen
(E)
Observed
Shannon
Library Method
SSLR
xGen
MDA_0.5h
MDA_1.5h
Nextera
(F)
Method
Virgaviridae
Unclassified
Unknown
MDA_0.5hvsNextera
MDA_1.5hvsNextera
SSLRvsNextera
xGenvsNextera
Phylum
Z-score
Library_Method
Nextera
MDA_0.5h
MDA_1.5h
SSLR
xGen
Phylum
Uroviricota
Unclassified
Kitrinoviricota
Unknown
Deseq2 p.adj value
1
0.05
0.001
* p.adj < 0.05

**Fig. S8** | (A) Distribution of sequencing reads into the different taxonomic categories as viral, human, bacterial, fungi and unknown origin. To check the presence of non-viral DNA sequences, 50,000 random forward reads were evaluated according to their match to a range of viral, bacterial, and human reference genome and protein databases. No reads (in 50,000 reads) matched the 18S rRNA gene sequences in all the samples. (B) The ratio of viruses and unknown part of fecal samples prepared with different methods. (C) The Venn diagram indicates the presence of vOTUs in the different library preparation methods. (D) The overall alignment rate (%) of high quality contigs that mapped to viral ref database. (E) The effect of different library preparation methods on the viral overall-alpha diversity with the measurement of Observed and Shannon index. NS indicates not significant, one asterisk indicates a significant difference at $p < 0.05$ (t.test), two asterisk indicates a significant difference at $p < 0.01$ (t.test), three asterisk indicates a very significant difference at $p < 0.001$ (t.test). (F) Differential abundance analysis by comparing the tested library preparation methods on the distribution of viruses at family level (Nextera XT library method was used as reference). Hierarchical clustering was performed to group family (rows) and library preparation methos (columns) based on similarities in differential abundance signatures and was visualized via heatmap.

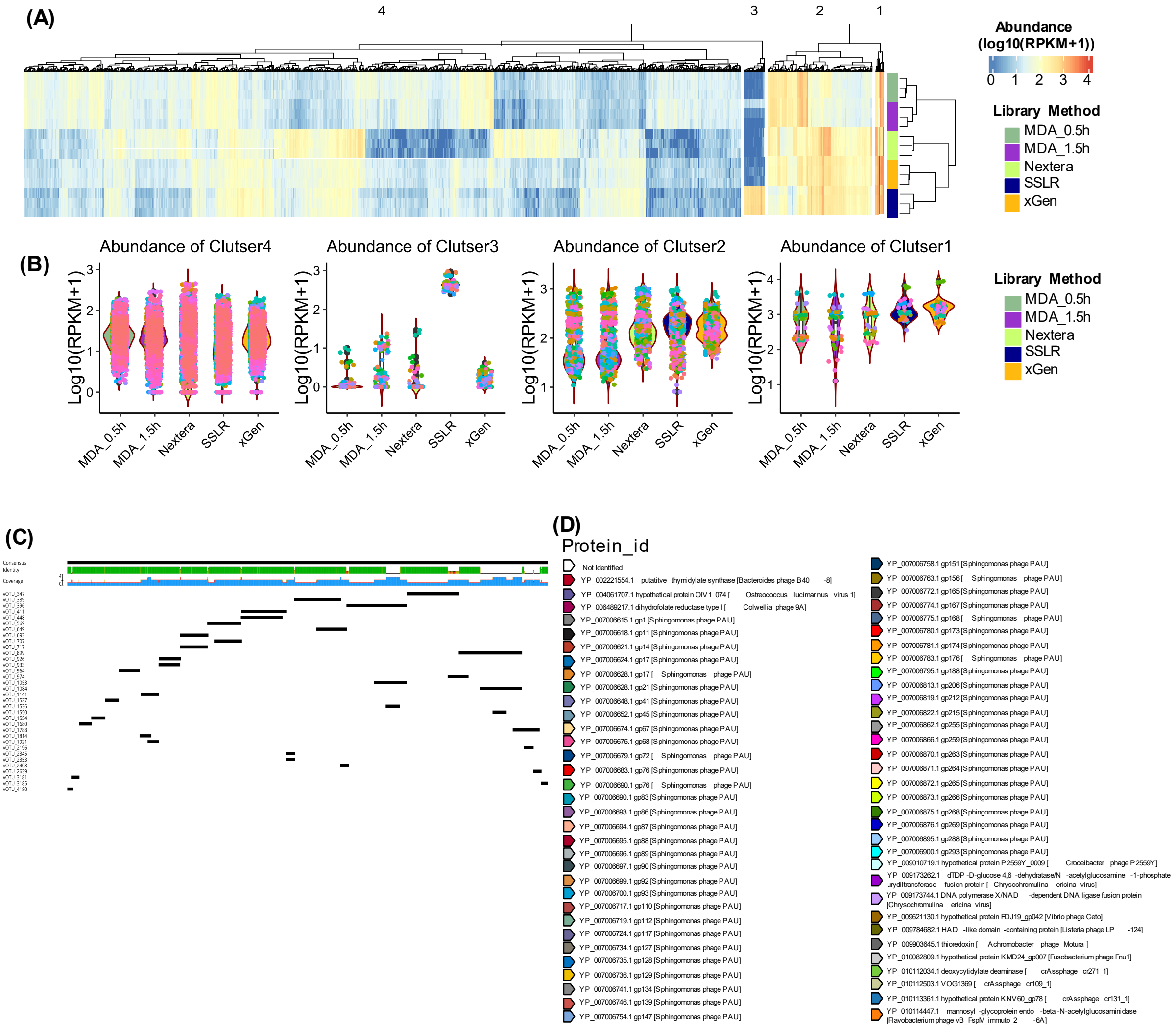


**Fig. S9** | (A) The heatmap shows the abundances (log10(RPKM+1)) of FV contigs, and the virome contigs were divided into 4 clusters. (B) The abundances of the 4 divided clusters. (C) 34 of high abundance (RPKM) viral contigs that were highly enriched in SSLR method that does not present in the rest of methods. Contig was piped through VirSorter 2 and DRAMv for viral identification and annotation. Multiple hits matched to Sphingomonas phage PAU (NC_019521.1). (D) Genomic maps of the open reading frames (ORFs) which are predicted by prodigal and then annotated by the blast from the NCBI protein database, the best hits were used for the annotation. Different colors indicate different annotated protein, directional boxes indicate ORFs in the respective orientation. Non-filled arrows indicate no protein hits, the rest of colored arrows are hits to majority of PAU proteins.

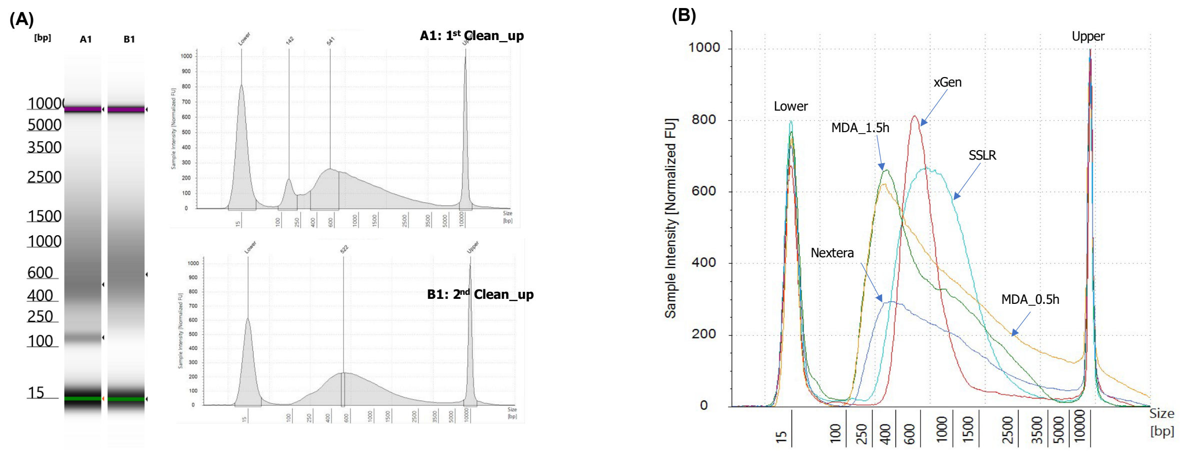


**Fig. S10** | Library size assessment by Agilent TapeStation 4200. (A) The extra clean-up step can remove the dimers that the first-round bead clean-up cannot remove. Fecal virome library prepared by the SSLR and clean-up by 0.7x AMPure XP beads. (B) Different library methods result in different library size distribution. In general, both MDA and Nextera XT tend to have shorter library size compared to single-stranded methods (SSLR and xGen).
