## Supplemental file1_Virome library workflow for "A single strand-based library preparation method for unbiased virome characterization"

**Protocol for metavirome sequencing with single-stranded library (SSLR) preparation**

### Introduction

We are here presenting a detailed workflow for virome library preparation with our lab-customed single-stranded library (SSLR). This workflow includes **7 FLEXIBLE** steps, virome isolation and extraction, mock community (positive control) preparation, reverse transcription with the purpose of dsRNA denatures, genome fragmentation, ligation with customed designed adaptors and library preparation for sequencing. Detailed information can be found on the next page.

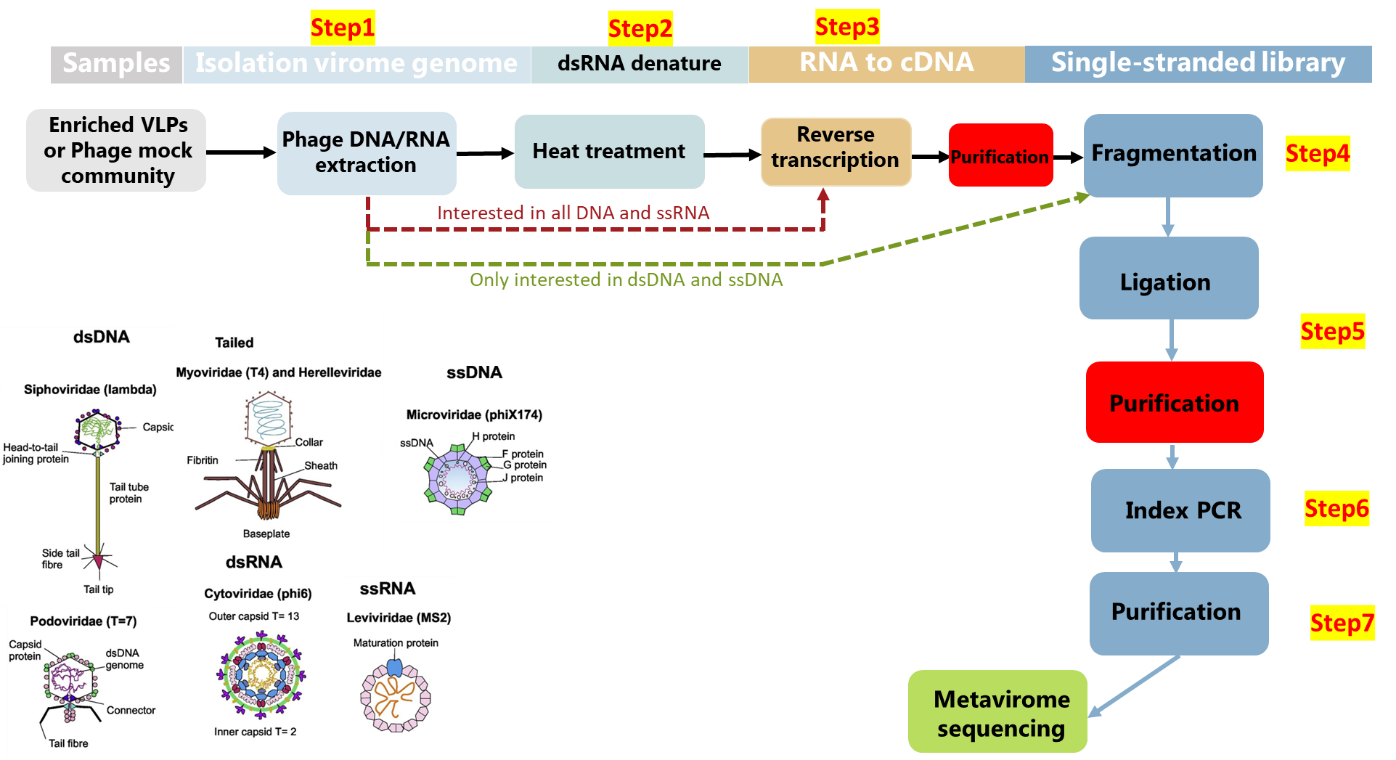

### Step1: Virome isolation and extraction

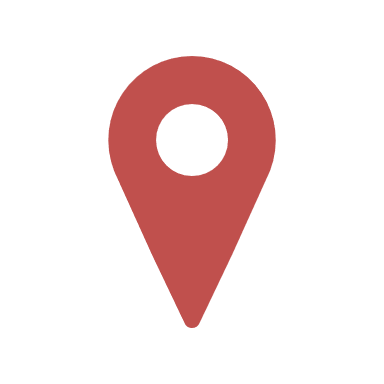

**Where	Timing: 180 min**

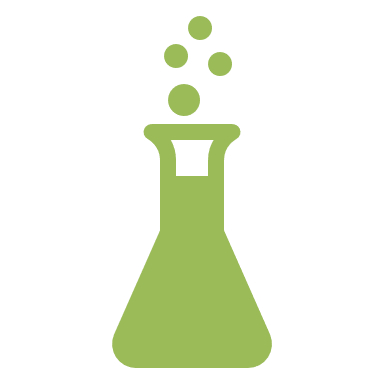
Fecal lab

**Reagents/kits**

SM buffer: lab-prepared according to <https://cshprotocols.cshlp.org/content/2006/1/pdb.rec8111.full?text_only=true>

Pierce™ Universal Nuclease for Cell Lysis: #88701

QIAamp Viral RNA Mini Kit (250): # 52906

**Equipment**

Centrifuge, 37°C incubator

**Procedure**

**Nuclease treatment and extraction**

1. Make aliquots of 140 µL of each of enriched sample.

2. Add 1 µL of 100-time diluted nuclease (**check the stock for the dilution in SM buffer**) to each sample and let them incubate at approx 30 min at 37°C. per. sample. Vortex between each sample

3. Consider increasing the incubation time if you expect a lot of external DNA.

4. Immediately after adding 540 µL AVL buffer to inactivate nucleases.

5. Mix the mixture by pulse vortexing.

6. Incubate at room temperature for 10 min.

7. Change gloves.

8. Briefly centrifuge the mixture with a microcentrifuge.

9. Add 560 μL of absolute ethanol (96%) to the sample mixture.

10. **CRITICAL STEP**: Mix very well by pulse-vortex.

11. Briefly centrifuge the mixture with a microcentrifuge.

12. Add 630 μL of the sample mixture to the spin column.

13. **CRITICAL STEP**: Do not touch the column rim with the pipet.

14. Centrifuge at 6000 × g for 1 min, change the collection tube and repeat steps 12 to load all the extracts.

15. Add 500 μL AW1 buffer to the spin column the following:

16. Centrifuge at 6000 × g for 1 min, change the collection tube.

17. Add 500 μL AW2 buffer to the spin column.

18. Centrifuge at 20,000 × g for 3 min, change the collection tube.

19. Centrifuge at 20,000 × g for 1 min. Place the spin column in a low-binding RNAse-free 1.5 mL tube.

20. Add 30 μL of AVE (elution buffer) to the spin column and incubate for 1 minute.

21. **CRITICAL STEP**: Pipet the AVE buffer directly onto the filter membrane without touching it with the pipet tip.

22. Centrifuge at 6000 × g for 1 min to collect the filtrates.

**PAUSE POINT** Send for next step or store viral DNA/RNA at –80°C.

**CAUTION**

- **NOT** interested in **RNA**??? **THEN GO STEP4**.
- Interested in **DNA** and **ssRNA**, **THEN GO STEP3**.
- Interested in DNA, dsRNA and ssRNA, **THEN GO STEP2**
- Remember to include a Positive (Mock from extraction) and Negative (H_2_O from extraction) controls for each extraction.

### Step2: Heat treatment

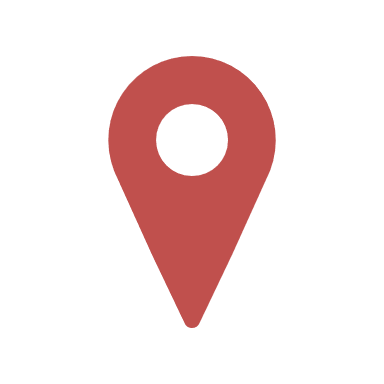

**Where	Timing: 3 min**

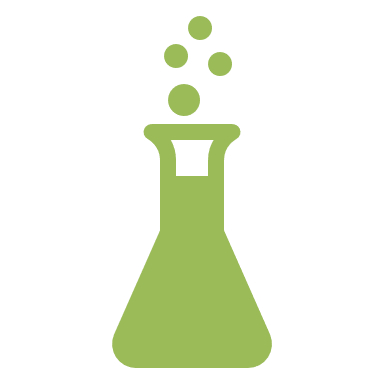
Clean-lab

**Reagents**

No reagents needed

**Equipment**

UV-beach, Thermocycler, Microcentrifuge

**Procedure**

1. Preheated ThermoCycle machine to 95°C and sterilized PCR tubes or plates depending on the how many samples are used for reverse transcription.

2. Add 16 μL of extracted virome in PCR tubes or plates, prepare on ice.

3. Briefly centrifuge the mixture with a microcentrifuge.

4. Put tubes or plates for heat treatment for 3 min.

5. **CRITICAL STEP**: transfer samples to ice immediately after 3 min.

6. Ready for reverse transcription.

**CAUTION**

- If you would like to look at the RNA in your samples, it is better to finish extraction, heat treatment and reverse transcription **on the same day**.
- Pipette, filtered pipette tips and workbench need to be sterilized by UV lamp for 20 minutes prior work.
- Remember to include a Positive (Mock from extraction), and Negative (H_2_O from extraction) controls for the reaction.

### Step3: Reverse transcription (RT)

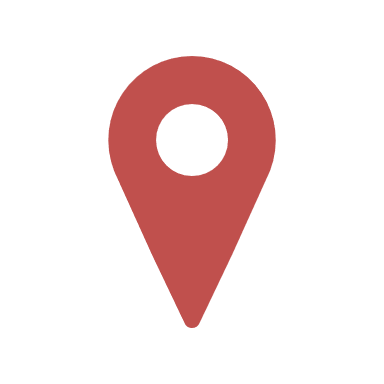

**Where	Timing: 25 min**

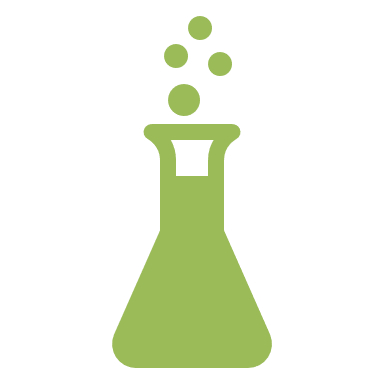
Clean-lab

**Reagents/kits**

SuperScript™ IV VILO™ Master Mix: #11756050

AMPure XP beads, 60 mL: #A63881

TE, pH 8.0, RNase-free: #AM9849

**Equipment**

Thermocycler, Microcentrifuge, UV workbench

**Procedure**

1. Transfer 16 μL of extracted or heat-treated virome DNA/RNA to a clean PCR plate.

2. Add 4 μL SuperScript™ IV VILO™ Master Mix to each sample and mix thoroughly, spin them down.

3. Carefully place the PCR plate into the ThermoCycle machine and select the following program:

| Temperature profile | | |
| --- | --- | --- |
| 25°C | 10 min | x 1 cycle |
| 50°C | 10 min |  |
| 85°C | 5 min |  |
| 4°C | ∞ |  |

4. Purified with 1X AMPure XP beads (25 μL /25 μL PCR reaction) and eluted with 20 μL Tris buffer (10 mM, pH8.0) according to the purification procedures from **Step7**.

**PAUSE POINT**: samples can be stored in fridge or freezer for downstream preparations.

**CAUTION**

- Pipette, filtered pipette tips and workbench need to be sterilized by UV lamp for 20 min prior work.
- During preparation, reagents and PCR plate must be on ice/cold block. Reagents are stored at –20°C.
- Spin down reagents before using them. Avoid vortexing enzymes.
- Remember to include a Positive (Mock from extraction), Negative (H_2_O from extraction) & Blank (H_2_O for the RT) controls for the reaction.

### Step4: Fragmentation

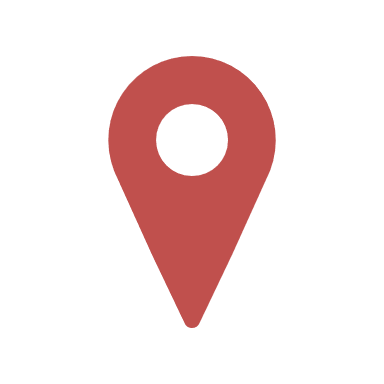

**Where	Timing: 15 min**

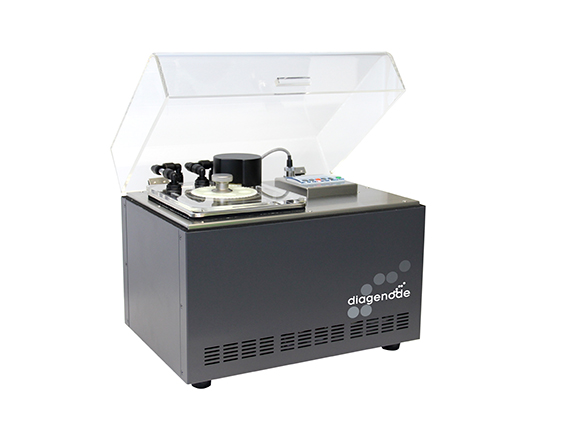

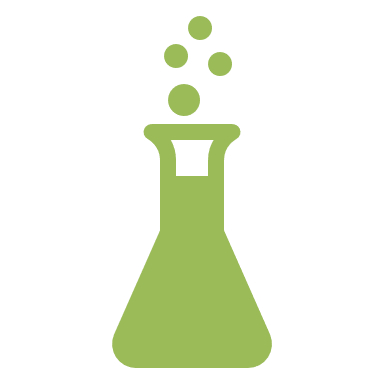
Basement (using ID card and key for access)

**Reagents**

No reagent needed

**Equipment**

Bioruptor® Pico sonication device (#B01060010), Microcentrifuge

**Procedure**

1. **CRITICAL STEP**: Pre-cooled (4°C) the Biorupter and holder at least 30 min.

2. While waiting for cooling down, transfer genomic DNA or reverse-transcription products to a new, clean Bioruptor tubes and briefly spin down to make sure all the liquid in the bottom of tubes.

3. Put samples on the ice for at least 10 min.

4. **CRITICAL STEP**: Set the sonication parameter as following: **15s ON and 90s OFF, using 8 cycles**

5. **CRITICAL STEP**: Spin down the samples and carefully load the Bioruptor tubes to sonicater holder and close carefully, make sure there is no liquid at the side of tubes.

6. **CRITICAL STEP**: Put the tube holder into the sonication chamber and close the lids. 12 samples can be done each time.

7. Spin down the tubes after shearing.

8. The fragmented product is ready for ligation.

**PAUSE POINT**: samples can be stored in fridge or freezer for downstream preparations.

**CAUTION**

- **Pregnant women** should not be stay away from the running machine.
- **DO NOT** turn on the instrument without water
- **Distilled water** should be used to fill the tank.
- **Always keep 12 tubes** for each run to a successful shearing.

### Step5: Ligation

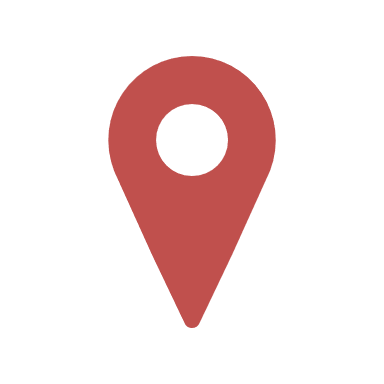

**Where	Timing: 60 min**

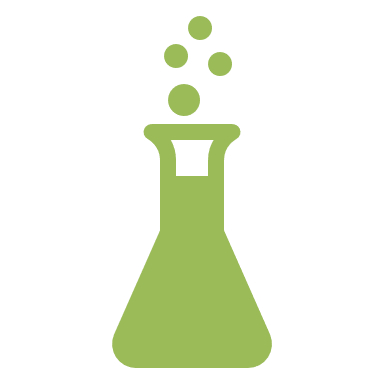
Clean-lab

**Reagents**

ET SSB (500 µg/mL): #M2401S

T4 Polynucleotide Kinase (10 U/μL): #EK0031

T4 DNA Ligase (5 U/μL): #EL0012

T4 DNA Ligase Buffer (10X) with 50% PEG-4000: #B69

Forward/reverse adapters: Lab designed and produced by IDT

AMPure XP beads, 60 mL: #A63881

**Equipment**

Thermocycler, Microcentrifuge

**Procedure**

- **Denature**

1. According to the number of samples (Include extra 3 for pipetting errors), prepare an ET SSB dilution in a clean tube. In the table below is calculated for a whole plate of 96 wells (Master Mix per 100 samples): diluted ET SSB to 5 ng/µL (500 µg /mL in the stock solution) with Tris and then add same volume of Tris, for example 1 µL ET SSB in 100 µL tris buffer. Mix them, spin them down and place on ice.

| **CAUTION**  **On ice** | **Reagent** | **Per 1 sample µL** | **Per 100 samples µL** |
| --- | --- | --- | --- |
|  | Fragmented DNA | 20 | - |
|  | ET SSB dilution (10 ng) | 2 | 100 |
|  | **Total denaturation reaction volume** | **22** |  |

2. Add 20 µL of fragmented DNA to each well.

3. Add 2 µL ET SSB solution, pipette several times and spin down.

4. Transfer to ThermoCycle and denature for 3 min at 95°C.

5. Cooling down on ice immediately after heating and set on ice for at least 5 min.

- **Ligation**

1. Spin down the heat denatured sample with a microcentrifuge.

2. According to the number of samples (Include extra 3 for pipetting errors), prepare a Master Mix containing per sample. In the table below is calculated for a whole plate of 96 wells (Master Mix per 100 samples). Mix them, spin them down and place on ice.

**CRITICAL STEP** for prepare master mix:

- Make sure the T4 ligase buffer and PEG are dissolve and mix thoroughly.
- **Add the needed volume of PEG-4000 (sticky, do it slowly), ligase buffer and PNK first and mix thoroughly, then add adapters and mix again, ligase should be added lastly during cooling down the samples and mix thoroughly without vortexing**.

3. Add 28 μL of ligation master mix to denatured samples, mix thoroughly with pipette, spin briefly.

4. Transfer to ThermoCycle for 60 min ligation at 37°C

|  | **95 ℃ for 3min, cool down on ice immediately for at least 5 min** | | |
| --- | --- | --- | --- |
|  | Reagents |  |  |
| **CAUTION**  **On ice** | PEG-4000 (50%) | 10 | 1000 |
|  | H_2_O | 8 | 800 |
|  | T4 ligase buffer (10X final) | 5 | 500 |
|  | T4 PNK (10,000 units/mL) | 1 | 100 |
|  | Add the above reagents one by one and then **mix for the 1^st^ time** | | |
|  | Forward adapter (1 pmol) | 1 | 100 |
|  | Reserve adapter (1 pmol) | 1 | 100 |
|  | Add the above adapters one by one and then **mix for the 2^nd^ time** | | |
|  | T4 DNA ligase (400,000 units/mL) | 2 | 200 |
|  | Add the above reagents one by one and then **by finger flicking to mix** | | |
|  | **Final volume** | **50** |  |
|  | **37 ℃ for 60 min (set the lid-heating off)** | | |

5. **Purified with 1X AMPure XP beads (25 μL /25 μL PCR reaction) and eluted with 22 μL MiliQ H_2_O according to the purification procedures from Step7.**

**PAUSE POINT** Store in fridge for short-time or freezer until beads clean-up.

**CAUTION**

- Ensure that you have booked a ThermoCycle and preheated it to 95°C.
- Pipette, filtered pipette tips and workbench need to be sterilized by UV lamp for 20 min prior work.
- During preparation, reagents and PCR plate must be on ice/cold block. Reagents are stored at –20°C.
- Spin down reagents before using them. Avoid vortexing enzymes.
- Remember to include a Positive (Mock from extraction), Negative (H_2_O from extraction) & Blank (H_2_O for the ligation) controls for the reaction.

### Step6: Index PCR

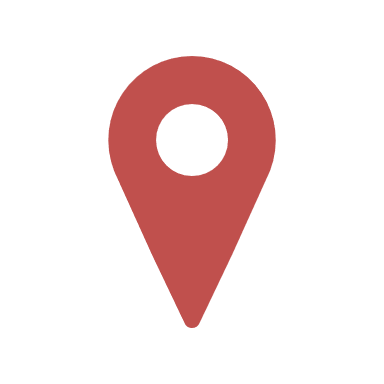
**Where	Timing: 30 min**

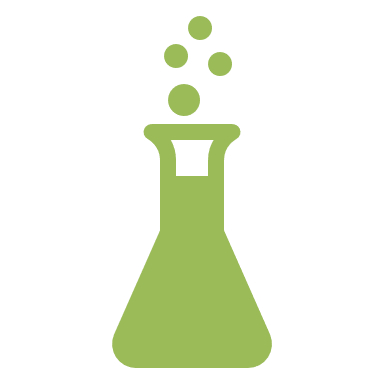
Seq-lab

**Reagents**

AccuPrime™ Taq DNA Polymerase System: #12339016

Nextera XT Index Kit v2 Sets: #FC-131-2001, 2002, 2003, 2004

**Equipment**

Thermocycler, Minicentrifuge

**Procedure**

1. Take the index primer plate from freezer to fridge prior to the start.

2. According to the number of samples (Include extra 3 for pipetting errors), prepare a Master Mix containing per sample. In the table below is calculated for a whole plate of 96 wells (Master Mix per 100 samples):

| **Reagent** | **Per 1 sample μL** | **Per 100 samples μL** |
| --- | --- | --- |
| Purified ligated DNA | 21.1 |  |
| 10× AccuPrime buffer | 2.5 | 250 |
| AccuPrime DNA polymerase | 0.4 | 40 |
| Primer P5 (i5) | 1 |  |
| Primer P7 (i7) |  |  |
| **Final volume** | **25** |  |
| The total volume for each well is 25 μL. |  |  |

3. In a clean EP tube, transfer the needed volume of 10× AccuPrime buffer and AccuPrime DNA polymerase from the stock solution, gently mix and spin down.

4. Transfer 2.9 μL of Master mix to each sample.

5. Transfer 1 μL of Primer mix to each sample.

6. Carefully mix them and spin them down.

7. Carefully place the PCR plate into the ThermoCycle machine and select the following program:

| Temperature profile | | |
| --- | --- | --- |
| 95°C | 2min |  |
| 95°C | 15s | x 20 cycles |
| 57°C | 30s |  |
| 68°C | 30s |  |
| 4°C | ∞ |  |

8. Purified with 0.8X AMPure XP beads (20 μL /25 μL PCR reaction) and eluted with 22 μL MiliQ H2O according to the purification procedures from **Step7.**

**PAUSE POINT** Store in fridge for short-time or freezer until beads clean-up.

**CAUTION**

- During the preparation of index PCR, reagents and PCR plate must be on ice/cold block. Lead opening while on cold block. Reagents are stored at –20°C.
- Gently mix and spin down reagents before using them. Avoid vortexing enzymes. 
- **WARNIN!!!** Change pipette tips and PCR Leads during work process!

### Step7: AMPure beads clean up and library quality check

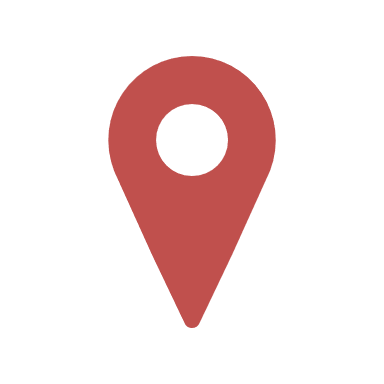

**Where	Timing: 30 min**

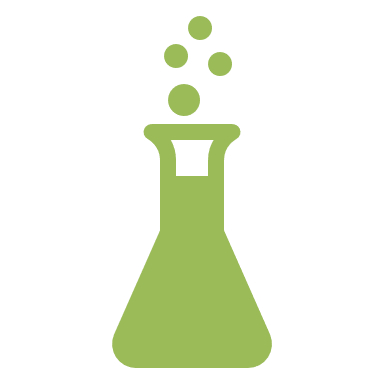
Seq-lab

**Reagents**

AMPure XP beads, 60 mL: #A63881

Qubit 1X dsDNA HS Assay Kit: #Q33231

High Sensitivity D5000 ScreenTape Assay: #5067-5593

**Equipments/kits**

HulaMixture, Invitrogen™ Qubit™ 4 Fluorometer, TapeSation4200 with high-sensitive D5000 Screen Tape (#5067-5592)

**Procedure**

1. Place AMPure XP beads into the Hula mixer to resuspend the beads and to equilibrate to room temperature for 15 min.

2. Label an Eppendorf tube or PCR plate with your sample id and transfer the 25 μl of PCR product.

3. Transfer 20 μL (!!! 0.8X) of Beads solution to each PCR product (25 μL) and mix with 100 μL pipette tips (10 times up and down), resulting 45 μL mixture.

4. Incubate for 5 minutes at room temperature.

5. Place the tube or plate with the mixture into magnetic rack for 2-4 min, until liquid is clear.

6. Carefully remove 40 μL of the liquid/supernatant with 100 μL pipette tips and discard it, keep 5 μL without disturbing the beads-pellet.

7. Wash beads-pellet with 175 μL freshly prepared 80 % EtOH by gently dispensing it over the beads with 200 μL pipette tips. Let it rest for 30 s and then remove the liquid.

8. Repeat washing step 7.

9. Spin the tudes or plates in a minicentrifuge for 30 s to collect all the residual liquid.

10. Put the tubes or plates on a magnetic rack and remove the excess with 10 μL pipette tips.

11. Air dry for approximately 30 s to evaporate the EtOH.

12. Remove the tube or plate from magnetic rack. Add 16 μL of nuclease free water and mix with pipette (>10 times up and down) to resuspend the beads-pellet. Incubate for 2 min. at room temperature.

13. Spin down for 30 s if there is liquid on the wall of tubes or well, place tube or plate back on the magnetic rack and wait for 3 min until the liquid clears.

14. Transfer 15 μL of the liquid/supernatant to a new, labeled Eppendorf tube or a new plate.

15. Qubit measurement of cleaned PCR products and pool 10 ng per sample to prepare a pooled library for library quality check with TapeStation following their standard protocols.

**PAUSE POINT** Send for sequencing or store in fridge for short-time or –80°C for long-time.
