## Supplemental file2_Adapters design for "A single strand-based library preparation method for unbiased virome characterization"

Additional file 2: SSLR adapter design

Forward adapter (P5):

/5AmMC6/TCGTCGGCAGCGTCAGATGTGTATAAGAGACAG

/3AmMO/AGCAGCCGTCGCAGTCTACACATATTCTCTGTCNNNNNNN/6CMmA5/

Reverse adapter (P7):

/5Phos/CTGTCTCTTATACACATCTCCGAGCCCACGAGAC/3AmMO/

/3AmMO/NNNNNNNGACAGAGAATATGTGTAGAGGCTCGGGTGCTCTG/5AmMC6/

3AmMO, 5AmMC6: modification and block parts

5Phos: 5’ Phosphate groups

Nucleotides in yellow background are the Illumina Paired-End Adapter Sequences
