## Supplemental file3_Data analysis for "A single strand-based library preparation method for unbiased virome characterization"

Additional file 3: Metavirome sequencing and data pre-processing

**Preprocessing of raw sequencing data of mock samples**

The mock metavirome generates a median of 2,483,360 reads with 150 bp paired-end sequencing (range, 165,994 - 12,192,992 reads, **Supplementary Table S8**). All the raw sequencing reads from Nextera XT, MDA_0.5h, MDA_1.5h and SSLR were trimmed from adaptors and barcodes and the high-quality sequences (>95% quality) using Trimmomatic v0.35 ^1^, with using NexteraPE-PE and a minimum size of 50nt were retained for further analysis. All the adaptors of raw reads from xGen library were removed by Trimmomatic and then the low complexity adaptase tail from first and last 8 nucleotides of each read were trimmed. After trimming, all the reads were subjected to within-sample de-novo assembly-only using metaSpades v3.15.1 ^2^, and the contigs with a minimum length of 2,200 nt were retained. Contigs generated from all samples were pooled and de-replicated at 90% identity using BBMap tool (dedupe.sh) ^3^. Prediction of viral contigs/genomes was carried out using VirSorter2 ^4^ (“full” categories|dsDNAphage, ssDNA, RNA, Lavidaviridae, nucleocytoplasmic large DNA viruses (NCLDV)|viral quality≥0.66), vibrant ^5^ (High-quality|Complete), and checkv ^6^ (High-quality|Complete). Taxonomy of mock community was inferred by blasting viral ORF against a customized phage mock taxonomy database (**Supplementary file6**) retrieved from NCBI and the Lowest Common Ancestor (LCA) for every contig was estimated based on a minimum e-value of 10e-5. Following assembly, quality control, and annotations, reads from all samples were mapped against the viral (high-quality) contigs (vOTUs) using the bowtie2 (version 2.2.5, default parameters) ^7^ and a contingency-table of reads per Kbp of contig sequence per million reads sample (RPKM) was generated, here defined as vOTU-table. Code describing this pipeline can be accessed on github: github.com/jcame/virome_analysis-FOOD. Genome assembly metrics before and after quality check were generated using QUAST (v.4.6.3) with default parameter ^8^. The coverage of each genome was generated by converting sam file of Bowtie2 alignment using samtools.

**Estimation of sequencing depths and error rates**

To estimate the sequencing depth of each phage genome from all mocks in the present study, SAM files generated by bowtie2 contigs mapping back to clean reads to BAM format using Samtools (v 1.9) Mapping percentage was calculated during Bowtie2 alignment and visualized by ggplot2.

To test whether heat and DMSO treatment cause any effect on sequencing fidelity during SSLR, we used two approaches to estimate sequencing error rates according to the previous method ^9^. R package ShadowRegression (v1.18) was used to calculate the error rate from each treatment, and then evaluated for differences using robust linear regressions ^10^. The detailed script can be found from: https://github.com/awilcox83/dsRNA-sequencing.

**Data pre-processing of metavirome of human fecal sequencing**

The fecal metavirome generates a median of 4,681,524 reads with 150 bp paired-end sequencing (range, 2,238,562 - 6,881,604 reads, **Supplementary Table S8**). The raw reads were trimmed as described above and checked for the presence of Phi X174 using BBMap tool (bbduk.sh) before MetaSpades assembly ^3^. Random forward reads of 50,000 of each samples were subjected to the Kraken2 program to map human, archaea, bacteria, viral, plasmid, UniVec Core with a standard pre-built reference database for the estimation of the contaminations ^11^. All the assembled contigs from each sample were pooled together and quality check with checkv, vibrant and virsorter was carried out as described above to generate high-quality contigs. Following assembly, quality control, and annotations, reads from all samples were mapped against the viral (high-quality) contigs (vOTUs) using the bowtie2 (version 2.2.5, default parameters) ^7^ and a contingency-table of reads per Kbp of contig sequence per million reads sample (RPKM) was generated, here defined as vOTU-table.

**Fecal virome taxonomy**

Taxonomy of the 1388 high-quality fecal virome contigs was inferred by blasting viral ORF against Virus Orthologous groups (VOG) release 217 (VOG217) database ^12^, COPSAC infant viruses ^13^ and human gut archaeal viruses ^14^ and the Lowest Common Ancestor (LCA) for every contig was estimated based on a minimum e-value of 10e-5.

**Host prediction of 34 representative virome contigs**

The 34 representative contigs were subjected to iPHoP v1.3.2 (integrated Phage Host Prediction) ^15^ to predict their host at genus level with default parameters by using its “add_to_db” function to add the 145 reference genomes of Sphingomonas species retrieved from NCBI.

**Searching PAU phage from 4 virus databases**

Usearch10 ^16^ and MMseqs2 ^17^ were used to search the existence of PAU phage from 4 gut virome databases, namely GVD (gut virome database) ^18^, GPD (gut phage database) ^19^, MGV (metagenomic gut virus) ^20^ and high-confidence IMG/VR4.1 (Integrated Microbial Genomes / Viruses) ^21^ with e-value of 10e-5. The best hits (top 5) were listed in **Supplementary Table S9**.
